# Population-specific transcriptional-state remodeling of cortical and hypothalamic neurons in Alzheimer’s disease

**DOI:** 10.64898/2026.08.15.745006

**Authors:** Sara Sejer, Magnus Møller Pedersen, Josephine Marie Boldt Lundby, Noor De Jong, Dong Won Kim

## Abstract

Selective neuronal vulnerability shapes Alzheimer’s disease, but it is commonly inferred from changes in the relative representation of neuronal populations rather than from molecular changes within those populations. We mapped both dimensions across disease progression in amyloid, tau, and combined amyloid-tau mouse models, extended the analysis to cortex and hypothalamus in an independent tau model, and tested the resulting framework in human Alzheimer’s disease.

Populations with similar reductions in representation showed markedly different degrees of molecular remodeling, while substantial state changes also occurred in populations whose representation remained close to wild type. Disease altered excitability, synaptic, and regulatory programs and reshaped the distribution of cellular states within matched neuronal identities.

A particularly recurrent excitatory state linked CAMKK2-AMPK signaling with microtubule regulation. It expanded in glutamatergic populations early in the combined amyloid-tau model, but its direction was not fixed: the same state reversed with age in dentate granule-like neurons and was reduced in corticothalamic neurons of an independent tau model. Remodeling extended beyond cortex to hypothalamic Hdc-positive tuberomammillary neurons, which showed focused excitability-associated changes despite representation close to wild type.

In human Alzheimer’s disease, remodeling was strongest in several deep-layer and RORB-positive excitatory populations independently associated with vulnerability. The CAMKK2-AMPK-associated state was predominantly reduced rather than increased. An independently defined human depletion-associated neuronal identity instead overlapped other synaptic and calcium-signaling components of the same broader remodeling architecture.

Neuronal involvement in Alzheimer’s disease is therefore expressed as structured, population-specific remodeling of molecular state, extending across cortical and hypothalamic identities and only partly reflected in relative representation.

## Introduction

Selective neuronal vulnerability is an organizing feature of Alzheimer’s disease. Neurons exposed to comparable amyloid and tau pathology do not degenerate equally, and neuronal loss follows stereotyped anatomical and cell-type patterns that pathological burden alone cannot explain^1–3^. Single-cell and single-nucleus transcriptomics have made these differences tractable at scale, and a common inference has followed: neuronal populations depleted in disease are considered vulnerable, whereas populations preserved in relative representation are considered comparatively resilient^4–7^.

Studies based on relative neuronal representation have identified selectively vulnerable excitatory populations and characterized molecular features associated with their identity. For example, quantitative compositional analyses across typical and atypical AD recently identified a reproducibly depleted NRGN/BEX1 excitatory population shared across clinical variants and brain regions and enriched for genes involved in synaptic transmission and plasticity^8^. These findings raise a distinct question: are the molecular features that mark a neuronal population as depletion-prone the same programs that change within neurons that remain represented in disease? Relative representation can identify populations whose recovery changes representation, but it provides limited information about the molecular state of the cells recovered within those populations. Resolving these dimensions separately is therefore necessary to distinguish constitutive features associated with depletion from disease-associated states superimposed on neuronal identity.

Three lines of evidence argue that preserved relative representation alone is insufficient to establish neuronal resilience.

First, neuronal function can fail even when structural innervation remains intact. In amyloid precursor protein knock-in mice, dopaminergic fibre density in the lateral entorhinal cortex is statistically indistinguishable from that of wild-type mice. Despite this structural preservation, cue-evoked dopamine release fails and associative memory encoding in entorhinal neurons collapses; both deficits are restored by L-DOPA^9^. Thus, a neuronal population can be functionally compromised, yet reversibly so, while appearing resilient when assessed by abundance alone.

Second, extensive molecular remodeling occurs in neurons that remain present in the tissue, demonstrating that state change and cell loss are mechanistically separable. Spatial single-cell proteomics of tangle-bearing neurons in post-mortem AD cortex shows that neurons occupy a continuum of proteomic states scaling with tau burden, progressing toward synaptic disruption with little evidence of activated cell-death programs^10^. This separation is further supported by direct experimental perturbations of neuronal chromatin maintenance, where disruption of Polycomb repressive complex 2 induces extensive cell-state and identity alterations without obligatory neuronal loss^11^, while progressive epigenetic derepression drives neurodegenerative phenotypes long before overt cell depletion^12^. Molecular remodeling can be separable from neuronal death and appears to be far more widespread among surviving neurons than the overt activation of cell-death programs.

Third, threshold models of selective vulnerability predict exactly this ordering: neurons differ in the stress load their intrinsic physiology can absorb, so molecular state precedes cellular fate rather than reporting on it^13–15^. This logic is not specific to AD. In Huntington’s disease, temporally resolved single-nucleus atlases identify coordinated remodeling of stress, proteostasis, and synaptic modules, together with progressive erosion of neuronal identity in projection neurons, before the onset of overt depletion^16^.

A more mundane problem compounds the conceptual one. Abundance estimates derived from dissociated or nuclei-isolated tissue are not unbiased measurements of how many cells are in a brain. Recovery efficiency varies with cell type, tissue integrity, and disease state, and enzymatic dissociation itself induces stress and immediate-early transcriptional programs that can be mistaken for biology^17,18^. In degenerating tissue, where the populations of interest are the most fragile, this bias runs in the direction most likely to produce false confidence. Vulnerability inferred from what is missing rests on two assumptions at once: that loss is the readout, and that loss was measured correctly.

Testing whether vulnerability exists as a state rather than an outcome imposes four requirements. (i) Transcriptional programs must be defined independently of the disease, so that the result is not the tautology that disease-derived signatures are altered in disease. (ii) Measurement must be at per-cell resolution within matched neuronal identities, so that a shift in the mean can be distinguished from the emergence of a discrete high-state subpopulation, a broadening of the distribution, or convergence onto a single new state. (iii) Where possible, the transcriptional state should be examined

against an independent regulatory measurement, to ask whether remodeling is accompanied by corresponding changes in chromatin accessibility. (iv) The framework requires an anchor outside the model system, since a mouse-defined program not engaged in human disease is of limited interest.

We address these in turn. For (i), we used differentially expressed genes across cortical neurons to identify candidate Reactome pathways, then retained each selected pathway with its full curated gene membership rather than only the differentially expressed genes within it. Pathways were grouped into three families covering synaptic organization, adhesion and neurotransmission (S); neuronal excitability, ion handling, Ca² -associated and G-protein-coupled receptor signaling (E); and transcriptional regulation, stress-associated signaling and cell-state control (R), with each pathway kept as an individual gene set rather than collapsed into a family-level aggregate. This selection step identifies which biological processes are examined but it does not determine which neuronal population, disease stage, or direction of change any program shows, and it cannot produce the opposing behavior of programs within the same family. As a sensitivity analysis, the ranking of the most recurrent excitability-associated program was also evaluated against the complete Reactome pathway collection. The resulting definitions were then held constant across genotype, age, neuronal population, model, platform, and species, so that every comparison outside the discovery cohort, including the independent tau model, the chromatin data, and the human atlas, used gene sets with no contact with the data being tested. Independent support for this program selection comes from an unbiased brain-wide screen of neuronal susceptibility to pathogenic tau and α-synuclein, conducted independently of expression level, which identified synapse- and Ca² -homeostasis genes as the principal modifiers of tau toxicity^19^, nominating from outside our data two of the three families that dominate the remodeling we observe.

For (ii), we profiled cortical cells from control, APP;PS1 (AP), Tau4RΔK, and Tau4RΔK-AP mice at 6, 9, and 12 months by single-cell RNA sequencing, with female mice as the primary disease-progression cohort and 12-month males as a biological reference with a less pronounced composition phenotype^20^, and evaluated the full per-cell distribution of program activity within each annotated population rather than its mean alone. For (iii), we integrated matched cortical single-cell chromatin-accessibility profiles from the same cohort. For (iv), we projected the mouse-derived programs into a published prefrontal cortex single-nucleus atlas of approximately 2.3 million cells from 427 ROSMAP donors spanning non-AD, early-AD, and late-AD neuropathological stages, with donor sex retained for stratified analysis^5^. We additionally tested reproducibility in an independent 6-month cortical and hypothalamic single-nucleus cohort from TauP301S-AP mice, carrying a distinct P301S tau transgene^21^ and profiled by a different cellular capture strategy, separating shared vulnerability-associated programs from effects specific to one tau line or platform. Finally, we asked whether the most recurrent of these programs is experimentally modifiable, using a pharmacologically perturbed neuronal-like cellular system.

We operationalized vulnerability-associated neuronal involvement across three separately quantified dimensions: change in relative population representation, remodeling of transcriptional programs within matched neuronal identities, and alteration of within-population state distributions. Relative representation was not equated with absolute neuronal loss, and these dimensions were not combined into a single vulnerability score. This framework allowed us to distinguish populations with similar representation changes but different molecular states, as well as populations that underwent substantial transcriptional remodeling while remaining comparatively close to wild type (WT) representation.

Using this framework, we mapped neuronal representation and molecular state across disease progression in Tau4RΔK-AP mice and its parental APP;PS1 and Tau4RΔK lines, then asked whether the resulting organization recurred in a genetically distinct TauP301S-AP model, in human AD neurons, and in a neuronal-like perturbation system. State remodeling was only partially aligned with relative representation and was organized across recurrent excitability-associated, synaptic, and regulatory programs. Individual pathway-associated states varied in magnitude and direction across neuronal identity, disease stage, model, and species, indicating that recurrence was stronger at the level of broader biological architecture than at the level of a fixed transcriptional signature. Together, these analyses show that neuronal involvement in Alzheimer’s disease cannot be inferred from relative representation alone. Instead, it is reflected in population-specific molecular states whose magnitude, distributional form, and direction depend on cellular and disease context.

## Results

### Changes in relative neuronal representation identify population-level shifts but not the state of surviving neurons

We mapped neuronal composition across disease progression by scRNA-seq in female control, APP;PS1 (APP), Tau4RΔK, and Tau4RΔK crossed with APP;PS1 (Tau4RΔK-AP) mice at 6, 9, and 12 months (Fig. 1A; WT, APP, Tau4RΔK, and Tau4RΔK-AP in figures). After removing glia and ambiguous populations, neurons were assigned to broad excitatory and inhibitory classes (Fig. 1B) and to six neuronal populations selected for composition analysis (Fig. S1A; Table S1): IT excitatory neurons (Ex1_IT), deep-layer L5 excitatory neurons (Ex2_L5deep), L6 corticothalamic neurons (Ex3_L6CT), dentate granule-like excitatory neurons (Ex4_DG_like), CGE-derived Calb2 interneurons (In1_CGE_Calb2), and a mixed MGE/CGE interneuron population analyzed at combined resolution (In2_MGE_CGE_mix). Claustrum/endopiriform Car3-like neurons, D1 and D2 striatal spiny projection neurons, and Cajal-Retzius cells were annotated but excluded from composition analyses. Given potential cell-type differences in dissociation efficiency, neuronal composition is reported as relative representation rather than absolute abundance.

**Figure 1.**
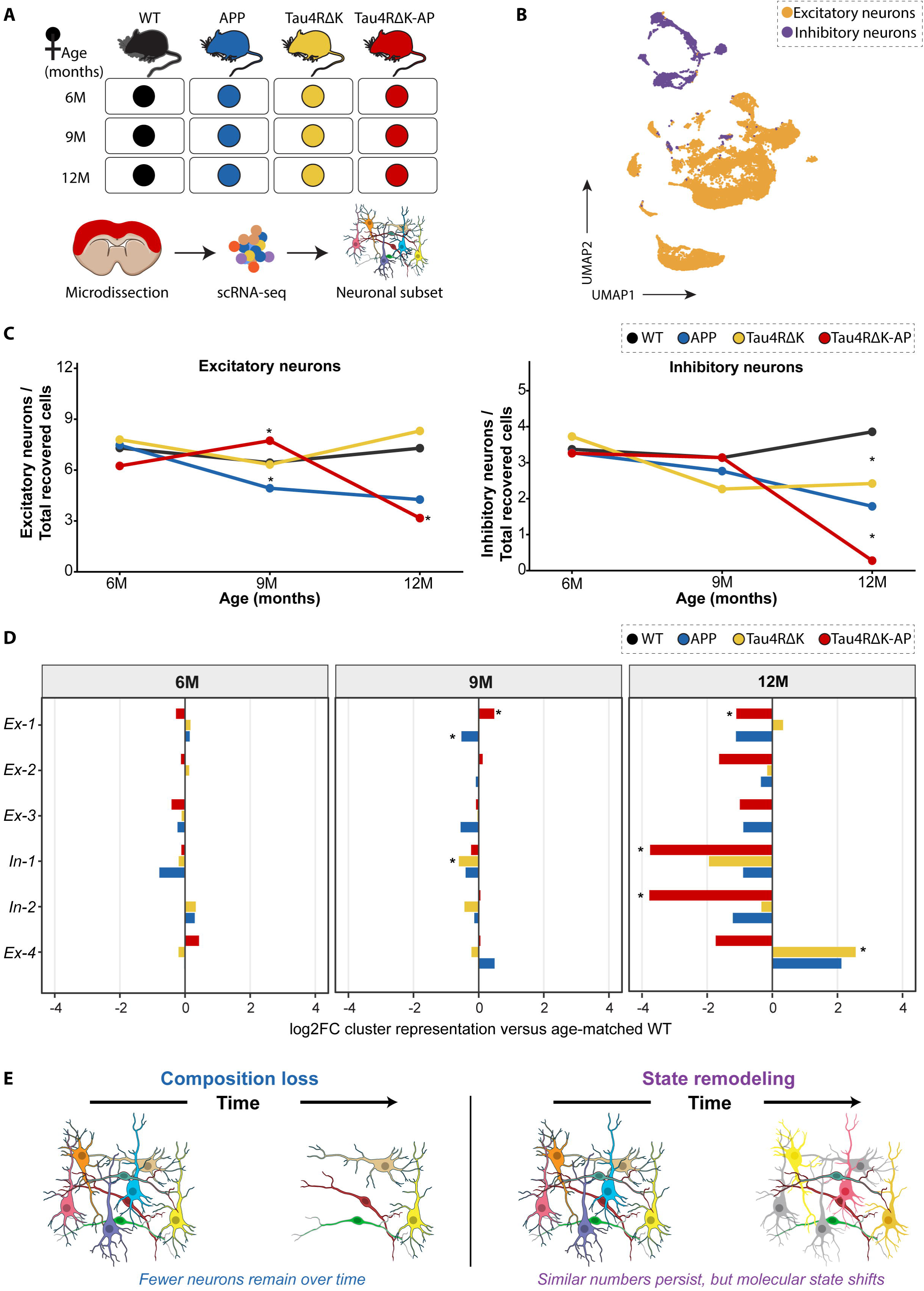
Changes in relative neuronal representation identify population-level shifts but do not define the state of surviving neurons. **(A)** Experimental design for the Tau4RΔK-AP disease-progression cohort. Female WT, APP;PS1, Tau4RΔK, and Tau4RΔK-AP mice were examined at 6, 9, and 12 months. Cortical tissue was profiled by single-cell RNA sequencing and neuronal populations were retained for downstream analysis. **(B)** Uniform manifold approximation and projection (UMAP) of cortical neurons colored by broad excitatory or inhibitory identity. **(C)** Relative representation of excitatory and inhibitory neurons as a fraction of all quality-controlled cortical cells recovered at each age. Points show genotype-level means of biological-sample representation, with lines connecting age-specific means. Genotype-associated differences were tested relative to age-matched WT using quasibinomial generalized linear models with biological sample as the unit of analysis. *p < 0.05. **(D)** Population-specific changes in relative neuronal representation across disease progression. Bars show the log2 fold change in representation of Ex1_IT, Ex2_L5deep, Ex3_L6CT, Ex4_DG_like, In1_CGE_Calb2, and In2_MGE_CGE_mix in APP;PS1, Tau4RΔK, and Tau4RΔK-AP relative to age-matched WT. Positive values indicate increased and negative values reduced relative representation. *p < 0.05 from the corresponding biological-sample-level quasibinomial models. **(E)** Conceptual distinction between relative neuronal representation and within-population molecular-state remodeling. A population can decrease in relative representation with either substantial or limited remodeling among recovered neurons, while a population with comparatively preserved representation can nevertheless undergo substantial molecular-state change. The schematic illustrates cross-sectional analytical dimensions and does not imply absolute neuronal loss or a fixed temporal progression.

Broad neuronal representation diverged by genotype and age (Fig. 1C). At 9 months, excitatory representation was lower than WT in APP mice (6.4% to 4.9%) and higher in Tau4RΔK-AP mice (6.4% to 7.7%). By 12 months, excitatory representation in Tau4RΔK-AP mice had fallen from 7.3% in WT to 3.2%. Inhibitory representation decreased in Tau4RΔK mice at 12 months (3.9% to 2.4%) and fell to 0.28% in Tau4RΔK-AP mice at 12 months, compared with age-matched WT. This approximately fourteen-fold difference also reduced the number of inhibitory neurons available for downstream analysis at that age. Thus, decreases in inhibitory representation were detected across three genotype-by-age comparisons, whereas excitatory changes were genotype- and age-specific.

These effects were not distributed evenly across neuronal identities (Fig. 1D). At 6 months, populations remained close to age-matched WT, with no nominally significant differences. At 9 months, Ex1_IT diverged in opposite directions between genotypes, decreasing in APP mice and increasing in Tau4RΔK-AP mice, while In1_CGE_Calb2 decreased in Tau4RΔK mice. At 12 months, In2_MGE_CGE_mix was reduced in all three disease genotypes, most strongly in Tau4RΔK-AP mice, where representation fell from 2.7% in WT to 0.19%. Ex1_IT was also reduced in Tau4RΔK-AP mice. Ex1_IT is a Rorb-like IT population broadly related to the human RORB-positive excitatory populations reported as selectively depleted in Alzheimer’s disease^7^. Ex4_DG_like increased in Tau4RΔK mice at 12 months. In1_CGE_Calb2 also showed a large, nominally significant reduction in Tau4RΔK-AP mice at 12 months, consistent with the strong negative change in relative representation shown in Fig. 1D. Thus, substantial representation changes at 12 months affected both excitatory and inhibitory identities, but their direction and magnitude remained population-specific.

A 12-month male cohort showed no statistically supported composition differences (Fig. S1B). Neither broad-class nor population-level comparisons reached nominal significance. Excitatory representation trended lower in all three disease genotypes, whereas total inhibitory representation was similar between WT and Tau4RΔK-AP mice. The 12-month male cohort therefore provides an additional biological context for examining molecular remodeling of the same neuronal identities, rather than a longitudinal comparison of sex-dependent trajectories.

None of these measurements reports the molecular state of the neurons that remain. A population reduced in representation may contain surviving neurons that are transcriptionally similar to WT or neurons that have undergone substantial remodeling. Conversely, a population maintained near WT representation may consist largely of neurons in an altered molecular state (Fig. 1E, Fig. S1C). Relative representation distinguishes depleted, comparatively stable, and enriched populations but is silent about the transcriptional state within them. We next asked whether disease reorganizes molecular programs within matched neuronal identities and whether the extent of this remodeling tracks changes in relative representation.

### Within matched neuronal identities, disease remodels synaptic, excitability-associated and regulatory states

To ask whether disease alters the molecular state of the neurons that remain, we used differentially expressed genes to nominate Reactome pathways, retained the full curated gene sets as scoring modules, and quantified module activity in individual neurons using AUCell (Fig. 2A; Tables S2 and S3). Modules were organized into three biological families: synaptic organization, adhesion and neurotransmission (S family; 24 modules); excitability, Ca² signaling and ion handling (E family; 22 modules); and transcriptional regulation and stress-associated signaling (R family; 25 modules) (Fig. S2A). The resulting discovery panel comprised 71 modules. These families provide an organizing framework rather than independent biological axes. Because Reactome is hierarchical, some modules are nested within broader pathways or span more than one family; these relationships are noted where they affect biological interpretation. Module names were retained from Reactome, including annotations originating from non-neuronal physiological contexts when the underlying gene content represented broadly expressed neuronal signaling machinery. Module activity was evaluated within six retained neuronal populations: Ex1_IT, Ex2_L5deep, Ex3_L6CT, Ex4_DG_like, In1_CGE_Calb2, and In2_MGE_CGE_mix.

**Figure 2.**
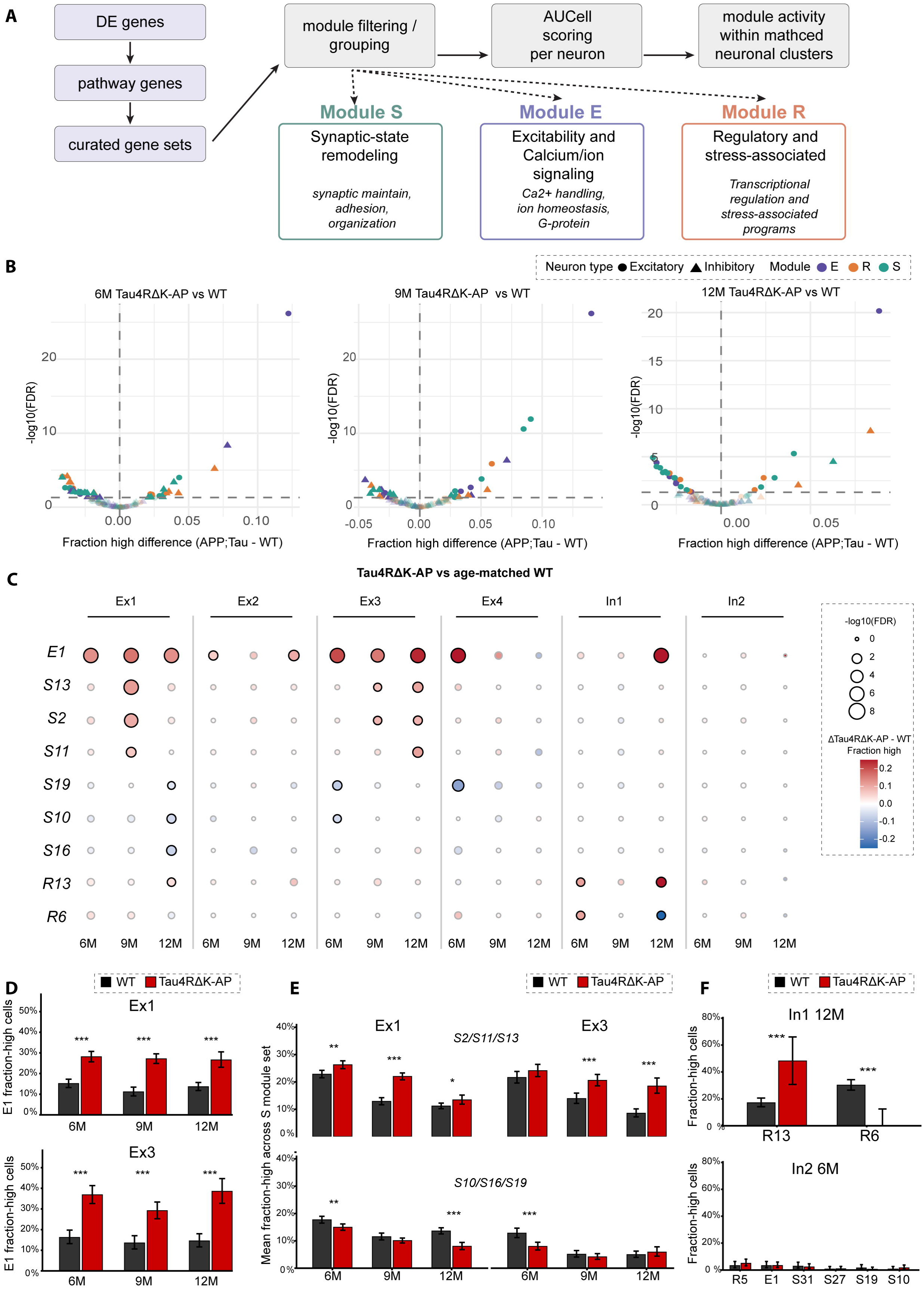
Tau4RΔK-AP-associated neuronal remodeling is organized along recurrent transcriptional-state axes within matched neuronal identities. **(A)** Overview of the transcriptional-state analysis. Disease-associated genes were used to nominate Reactome pathways, after which complete curated pathway gene sets were retained as scoring modules and quantified in individual neurons using AUCell. Modules were organized into synaptic-state (S), excitability and calcium/ion-signaling (E), and regulatory and stress-associated (R) families. **(B)** Global distribution of Tau4RΔK-AP-associated module-state changes at 6, 9, and 12 months relative to age-matched WT. Each point represents a module by neuronal-population comparison. The x-axis shows the Tau4RΔK-AP-minus-WT difference in the fraction of module-high cells (Δfraction-high), with positive values indicating increased and negative values reduced module-high occupancy. Point shape denotes excitatory or inhibitory identity and color denotes module family. The y-axis shows cell-level FDR from comparisons of module-high and non-high cellular frequencies and is used to characterize separation of the sampled cellular-state distributions. **(C)** Selected module-state changes across Ex1_IT, Ex2_L5deep, Ex3_L6CT, Ex4_DG_like, In1_CGE_Calb2, and In2_MGE_CGE_mix across age. Columns represent neuronal population by age combinations and rows represent selected E-, S-, and R-family modules. Circle color shows signed Δfraction-high, with red indicating increased and blue reduced occupancy in Tau4RΔK-AP. Circle size represents absolute effect magnitude, |Δfraction-high|. **(D)** Fraction of E1-high cells in Ex1_IT and Ex3_L6CT neurons from WT and Tau4RΔK-AP mice at 6, 9, and 12 months. E1 corresponds to the NMDAR-linked AMPK signaling program. Bars show pooled cellular fractions and error bars show 95% Wilson score intervals. **(E)** Module-high occupancy for selected postsynaptic and synaptic-state programs in Ex1_IT and Ex3_L6CT neurons. S2, S11, and S13 represent nested postsynaptic signaling programs, whereas S19, S10, and S16 represent receptor-gating and synaptic-adhesion-associated programs. Bars show pooled cellular fractions and error bars show 95% Wilson score intervals. **(F)** Selected regulatory and excitability-associated remodeling in inhibitory populations. Top, R13-high and R6-high fractions in In1_CGE_Calb2 at 12 months. Bottom, module-high fractions for selected E-, S-, and R-family programs in In2_MGE_CGE_mix at 6 months. Bars show pooled cellular fractions and error bars show 95% Wilson score intervals.

Within this discovery-informed panel, recurrent remodeling involved a restricted set of excitability-associated, synaptic, and regulatory programs (Fig. 2B). E1 was among the prominent positive effects, while selected synaptic and regulatory modules showed both positive and negative remodeling depending on neuronal identity and age.

### Excitability-associated remodeling emerges early in excitatory neuronal populations

Within the E-family, E1 was one of the most recurrent positive states in Tau4RΔK-AP neurons (Fig. 2B, C). E1 corresponds to the Reactome pathway *Activation of AMPK downstream of NMDARs*. At the better-sampled 6- and 9-month comparisons, E1 shifted positively across the excitatory populations examined while remaining close to WT in both inhibitory populations. Ex1_IT and Ex3_L6CT showed persistent increases across the progression series, whereas Ex2_L5deep showed smaller effects. Ex4_DG_like followed a distinct temporal pattern, with strong E1 expansion at 6 months followed by attenuation and reversal by 12 months. In1_CGE_Calb2 showed little E1 remodeling at 6 and 9 months but increased at 12 months, while In2_MGE_CGE_mix remained comparatively stable.

E1 occupies a distal position within the Reactome postsynaptic NMDAR hierarchy. The finalized scoring module contains *Camkk2*, AMPK subunit genes, and α- and β-tubulin genes represented in the curated NMDAR-AMPK-microtubule pathway; *Mapt* was excluded from the scoring module. E1 is interpreted here as a pathway-associated transcriptional state rather than as a direct measure of AMPK kinase activity. The underlying signaling architecture is nevertheless relevant to AD, as CAMKK2-AMPK signaling has been linked experimentally to Aβ-associated tau phosphorylation and synaptic dysfunction^22,23^. E1 remodeling was already evident at 6 months, when changes in relative representation were comparatively limited, but its subsequent direction varied with neuronal identity and age.

### Postsynaptic signaling and receptor-proximal programs remodel in opposing directions

Synaptic programs separated into opposing patterns (Fig. 2C, E; Fig. S3A). The nested postsynaptic modules S11 (Neurotransmitter receptors and postsynaptic signal transmission), S2 (Activation of NMDA receptors and postsynaptic events), and S13 (Post NMDA receptor activation events) generally increased, whereas the receptor-proximal S19 branch and the synaptic-adhesion modules S10 and S16 decreased in selected populations. Together, these opposing changes indicate a coordinated redistribution across the postsynaptic architecture rather than uniform activation or suppression of synaptic transcription.

In contrast, S19 (Unblocking of NMDA receptors, glutamate binding and activation), S10 (Neurexins and neuroligins), and S16 (Synaptic adhesion-like molecules) showed lower module-high fractions in several matched comparisons. The combined decrease was most evident in Ex1_IT at 6 and 12 months and in Ex3_L6CT at 6 months (Fig. 2E; Fig. S3A). S19 is the receptor-proximal sibling branch of S13 within S2, whereas S10 and S16 represent trans-synaptic adhesion programs outside the NMDAR hierarchy.

The resulting pattern was positionally organized. Downstream NMDAR signaling increased while the receptor-gating branch and selected trans-synaptic adhesion programs decreased within the same neuronal identities. These opposing changes are inconsistent with a uniform reduction in synaptic transcription and instead indicate reorganization of postsynaptic signaling relative to receptor-proximal and structural synaptic programs.

### Regulatory remodeling differs between inhibitory populations and disease stages

Regulatory-state remodeling was strongly restricted by neuronal identity and age (Fig. 2C, F). In In1_CGE_Calb2 neurons at 12 months, the R13-high fraction (Regulation of TP53 Activity through Methylation) increased from approximately 16% to 48%, whereas the R6-high fraction decreased from approximately 30% to near zero. R6 is annotated in Reactome as the Gastrin-CREB signalling pathway via PKC and MAPK and comprises PKC-, MAPK-, and CREB-associated signaling components. The opposing changes indicate redistribution between regulatory programs rather than uniform activation of the R family.

In contrast, In2_MGE_CGE_mix showed comparatively little remodeling across the selected module panel. At 6 months, the R5, E1, S31, S27, S19, and S10 module-high fractions were all low and showed only small genotype differences (Fig. 2F). Thus, the pronounced late regulatory remodeling observed in In1_CGE_Calb2 was not a shared feature of the two retained inhibitory populations.

Populations also differed markedly in remodeling breadth. Ex2_L5deep showed relatively restricted remodeling, dominated by E1, whereas Ex1_IT and Ex3_L6CT showed broader changes across excitability and synaptic programs (Fig. 2C). Ex4_DG_like was distinguished by a strong but transient E1 response, with maximal expansion at 6 months followed by loss and reversal of this response by 12 months. In2_MGE_CGE_mix showed comparatively little remodeling across the selected panel.

### Remodeling is supported at the biological-replicate level and is not explained by threshold choice or detection depth

At the biological-replicate level, we aggregated expression within each matched neuronal population per animal and compared pseudobulk module-gene enrichment with the Tau4RΔK-AP versus WT change in module-high fraction (Fig. S3B). Correspondence between the two measures was most evident for E-family programs, for which the largest increases in module-high fraction were associated with positive pseudobulk enrichment. S- and R-family programs were more dispersed, consistent with their bidirectional and population-dependent behavior. These analyses provide complementary biological-replicate support for the cell-state changes identified by AUCell.

The direction of remodeling was generally stable across alternative definitions of the module-high state (Fig. S2B, C). Using WT-derived 85th, 90th, and 95th percentile thresholds, denoted q85, q90, and q95, E-family comparisons retained the same direction in 18 of 18 comparisons, R-family comparisons in 40 of 54, and S-family comparisons in 81 of 108. Threshold choice affected effect magnitude and whether individual comparisons met the cell-level FDR criterion, but the principal directional patterns were preserved.

The principal excitatory effects were also not explained by transcript detection depth. In Ex1_IT and Ex3_L6CT, WT neurons showed slightly higher baseline median UMI counts and detected gene numbers than Tau4RΔK-AP neurons (Table S4), so the higher E1 state was observed despite lower transcript recovery in disease. E1 AUCell activity remained higher in Tau4RΔK-AP neurons in linear mixed-effects models that included UMI depth and detected gene number as covariates and animal as a random intercept. Selected E-, S-, and R-family effects also retained their direction and approximate magnitude after 1:1 nearest-neighbor matching on detected gene number and UMI count, with effects evaluated within population and age after matching. E1 additionally retained its disease-associated increase after fixed-depth UMI downsampling and recalculation of AUCell scores (Table S4). The bidirectional remodeling observed within the same neuronal populations provides a further control against a uniform detection effect: E1 and S13 increased while S19, S10, and S16 decreased (Fig. 2C, E; Fig. S3A).

### Tau4R***Δ***K and APP;PS1 contribute different components at different ages

The single-transgene lines resolved distinct components of the Tau4RΔK-AP remodeling profile across disease progression (Fig. S4A, B). Tau4RΔK mice showed E1 elevation in Ex1_IT from 6 months onward and in Ex3_L6CT from 9 months, together with bidirectional synaptic remodeling from the earliest age examined. Reduced S10, S16, and S19 states were evident in Ex1_IT and Ex3_L6CT across selected ages. Excitatory representation in Tau4RΔK mice remained close to WT across all three ages (Fig. 1C), providing a direct example of substantial within-population remodeling despite comparatively preserved representation of the excitatory compartment.

APP;PS1 mice followed a different course, with limited remodeling at 6 and 9 months and more prominent changes at 12 months. These later effects were concentrated in the nested S2/S11/S13 postsynaptic signaling program in Ex1_IT and Ex3_L6CT, with weaker involvement of the distal E1 branch (Fig. S4A).

The Tau4RΔK-AP profile was not reproduced in full by either parental genotype. Tau4RΔK mice showed early remodeling across excitability-associated and synaptic programs, including E1, whereas APP;PS1 mice showed more prominent later remodeling of the broader S2/S11/S13 postsynaptic hierarchy. The Tau4RΔK-AP profile was not reproduced in full by either parental genotype.

### The male Tau4R***Δ***K-AP cohort shows a smaller and distinct remodeling profile

The 12-month male Tau4RΔK-AP cohort showed fewer selected module changes than the female cohort (Fig. S4C). E1 increased in Ex1_IT, and S16 also increased in Ex1_IT. The strongest inhibitory effect was an increase in R13 in In1_CGE_Calb2. Other selected programs showed smaller population-specific shifts, whereas In2_MGE_CGE_mix remained comparatively close to WT across the displayed module panel. Thus, the male data did not reproduce the late female In1_CGE_Calb2 regulatory pattern or provide evidence for substantial E1 remodeling in In2_MGE_CGE_mix.

Broad inhibitory representation remained comparable between WT and Tau4RΔK-AP mice in the male cohort at this age (Fig. S1B). The presence of a strong S16 state change in In1_CGE_Calb2 despite preserved broad inhibitory representation provides an additional example of molecular remodeling without a corresponding broad-class representation change. Because the male cohort was sampled only at 12 months, these differences do not distinguish a sex-dependent trajectory from cohort-specific variation.

### Within-population state remodeling is not captured by relative neuronal representation

Having identified recurrent transcriptional-state changes within matched neuronal identities, we next asked how remodeling related to changes in relative representation. We considered population-level representation together with module-high state change and examined complete per-cell AUCell distributions in selected neuronal populations (Fig. 3).

**Figure 3.**
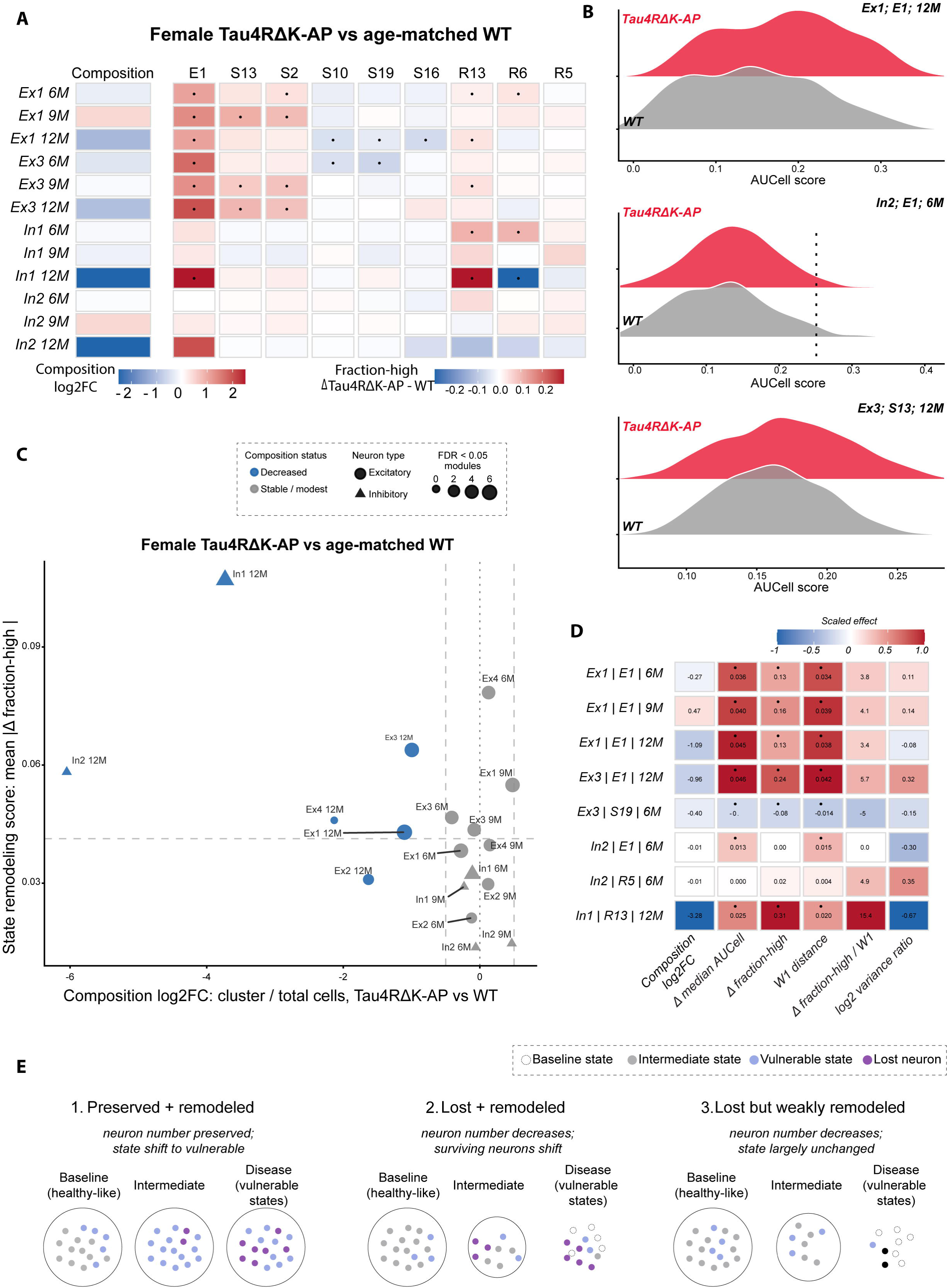
Within-population distributional remodeling dissociates from relative neuronal representation. **(A)** Joint visualization of relative neuronal representation and selected transcriptional-state effects across the female Tau4RΔK-AP progression cohort. The first column shows population-level representation log2 fold change relative to age-matched WT. Remaining columns show Tau4RΔK-AP-minus-WT Δfraction-high for selected E-, S-, and R-family modules. Red indicates positive and blue negative effects. **(B)** Representative per-cell AUCell-score distributions illustrating distinct forms of state remodeling. Distributions are shown for E1 in Ex1_IT at 12 months, E1 in In2_MGE_CGE_mix at 6 months, and S13 in Ex3_L6CT at 12 months. WT distributions are shown in gray and Tau4RΔK-AP distributions in red. The WT-derived q90 module-high threshold is indicated where shown. **(C)** Relationship between relative neuronal representation and aggregate transcriptional-state remodeling across population-age combinations. The x-axis shows population-level representation log2 fold change relative to age-matched WT and the y-axis shows mean absolute Δfraction-high across the selected module panel. Point shape denotes excitatory or inhibitory identity and point color denotes descriptive representation category. Vertical reference lines indicate the predefined representation boundaries at log2FC = -0.5 and +0.5. **(D)** Distributional characterization of selected neuronal population by module by age combinations. Columns show population-representation log2 fold change, change in median AUCell score, Δfraction-high, direction-signed Wasserstein-1 distance, and log2 variance ratio. Values are scaled within each metric for visualization, with red and blue indicating positive and negative directional effects. **(E)** Conceptual summary of three observed relationships between relative representation and molecular state: preserved or increased representation with substantial remodeling, reduced representation with substantial remodeling, and reduced representation with comparatively limited remodeling.

Changes in relative representation and within-population state remodeling followed distinct patterns across the female Tau4RΔK-AP progression cohort (Fig. 3A, C; Table S5). Ex1_IT and Ex3_L6CT showed recurrent remodeling across excitability-associated and synaptic programs, while their relative representation varied in magnitude and direction across age. In1_CGE_Calb2 showed comparatively limited remodeling at 6 and 9 months, followed by pronounced excitability-associated and regulatory remodeling at 12 months together with a large reduction in representation. In2_MGE_CGE_mix followed a different pattern, with relatively little remodeling across the selected module panel despite a marked reduction in representation at 12 months.

The clearest contrasting example occurred in Ex4_DG_like at 6 months. This population showed one of the largest aggregate remodeling scores in the dataset while its relative representation remained close to age-matched WT (Fig. 3C). Ex1_IT at 9 months provided a second example, combining substantial remodeling with increased rather than reduced representation. Thus, large state changes were not restricted to populations with reduced representation, and populations with similar representation outcomes could differ substantially in molecular-state remodeling. Relative representation and molecular state therefore provided non-redundant information about the same neuronal populations.

### Similar fraction-high changes can arise from different distributional rearrangements

Per-cell AUCell distributions showed that changes in molecular state were not captured completely by threshold occupancy alone (Fig. 3B; Fig. S5A). Selected programs showed several forms of redistribution. In Ex1_IT neurons at 12 months, the E1-associated distribution shifted toward higher scores, while Ex3_L6CT neurons showed a similar displacement for S13. In In2_MGE_CGE_mix neurons at 6 months, E1 showed a modest rightward displacement despite essentially unchanged q90 high-state occupancy. Median score and signed Wasserstein-1 distance increased, whereas the q90 fraction-high difference remained near zero (Fig. 3D). Threshold occupancy therefore captured only one aspect of remodeling and could differ from changes in the location or shape of the complete cellular-state distribution.

### Strong remodeling occurs across the full range of relative representation change

We next compared relative representation with an aggregate measure of module-state remodeling, defined as the mean absolute change in fraction-high across the selected modules for each population and age (Fig. 3C; Table S6). Remodeling occurred across a broad range of representation changes rather than being restricted to populations with reduced representation.

Ex4_DG_like at 6 months provided the clearest example of preserved representation with strong remodeling, showing one of the largest remodeling scores while remaining close to WT in relative representation. Ex1_IT at 9 months likewise combined increased representation with substantial remodeling. Among populations with reduced representation, In1_CGE_Calb2 at 12 months showed the largest remodeling score, whereas In2_MGE_CGE_mix at 12 months was also strongly reduced but showed a substantially smaller remodeling score. Ex3_L6CT at 12 months occupied an intermediate position, with reduced representation and strong state remodeling. Populations with similar qualitative representation outcomes could differ markedly in molecular-state remodeling, and substantial remodeling was not restricted to populations with reduced relative representation.

### Displacement, threshold occupancy, and distributional width are separable outcomes

Selected population-module combinations illustrated distinct forms of remodeling (Fig. 3D; Table S7). Ex1_IT E1 and Ex3_L6CT S13 showed concordant increases in central tendency, fraction-high occupancy, and signed Wasserstein-1 distance. By contrast, Ex3_L6CT S19 at 6 months shifted in the opposite direction, showing that different programs within the same neuronal identity could remodel oppositely. In2_MGE_CGE_mix at 6 months showed a more metric-dependent pattern, with modest displacement of the E1 distribution but essentially unchanged q90 occupancy. In1_CGE_Calb2 R13 at 12 months showed yet another configuration, combining increased high-state occupancy with pronounced narrowing of the distribution.

### Threshold sensitivity distinguishes stable directional effects from distribution-dependent effects

For selected population-module examples, threshold sensitivity was evaluated using WT-derived q80, q90, and q95 definitions (Fig. S5B; Table S8). Several major effects, including Ex1_IT E1, Ex3_L6CT S13, and the decreases in Ex3_L6CT S10 and S19, retained their direction across thresholds. In contrast, the small early effects in In2_MGE_CGE_mix were more threshold-dependent.

The late R13 and R6 effects in In1_CGE_Calb2 attenuated at q95, consistent with the distributional geometry observed in Fig. 3D. This sensitivity reflected where the disease and WT distributions crossed the selected threshold rather than reversal of the underlying state difference. Together, these analyses show that threshold-defined occupancy is useful for summarizing remodeling but should be interpreted together with the full score distribution.

### Three patterns of neuronal involvement

These analyses distinguish three descriptive patterns of neuronal involvement (Fig. 3E): populations with preserved or increased representation and substantial molecular remodeling, populations with reduced representation in which the remaining neurons are strongly remodeled, and populations with reduced representation but comparatively limited detectable remodeling. Ex4_DG_like at 6 months exemplified the first pattern, In1_CGE_Calb2 at 12 months the second, and In2_MGE_CGE_mix at 12 months the third.

These analyses separate three dimensions of neuronal involvement: relative population representation, the magnitude of molecular-state remodeling within an identity, and the distributional form of that remodeling. These dimensions showed incomplete correspondence across neuronal populations and disease stages, indicating that changes in population representation and molecular state capture distinct features of disease-associated neuronal remodeling.

### Early module state does not order later age-associated changes in relative representation

As an exploratory age-structured analysis, we asked whether early state remodeling ordered later age-associated changes in relative representation (Fig. S6; Table S9). Across the five populations with sufficient data, neither 6-month E1 remodeling nor the broader module panel showed a robust monotonic association with representation differences observed at later ages after multiple-testing correction. Thus, early transcriptional remodeling did not provide a simple ordering of subsequent age-associated shifts in neuronal representation. Given the limited number of populations and cross-sectional sampling, this analysis was used to assess association rather than longitudinal prediction.

### Chromatin accessibility at module-associated loci

To ask whether RNA-defined state remodeling was accompanied by changes in chromatin accessibility, we analyzed matched cortical scATAC-seq data (Fig. S7A; Tables S10, S11). The clearest directional correspondence occurred for the receptor-gating and adhesion modules S19, S10, and S16 in aEx3, broadly corresponding to RNA-defined Ex3_L6CT. Their GeneActivity changes matched the direction of the RNA-level effects (Fig. S7B). In contrast, the nested postsynaptic modules S13, S2, and S11 did not show corresponding increases in GeneActivity, and E1 likewise showed no consistent accessibility counterpart to its RNA-level remodeling (Fig. S7C). Regulatory modules showed heterogeneous GeneActivity changes across populations and ages, without a consistent cross-modal pattern (Fig. S7D).

Across matched population-module comparisons, RNA module-high changes and ATAC GeneActivity changes showed no global association (Fig. S7E). Cross-modal correspondence was therefore selective, with the clearest accessibility correlates observed for the decreasing receptor-gating and adhesion programs.

Together, these analyses separate three features of neuronal involvement: relative population representation, the magnitude of within-identity transcriptional remodeling, and the distributional form of that remodeling. Populations with similar representation changes could occupy substantially different molecular states, while strong remodeling could occur in populations whose representation remained close to WT. Relative representation therefore does not recover the molecular state of neurons within a matched population.

### Mouse-derived neuronal-state programs show selective and stage-dependent engagement in human Alzheimer’s disease neurons

To ask whether the neuronal-state architecture defined in mouse is engaged in human disease, we scored human orthologs of the fixed S-, E-, and R-family modules across annotated neuronal populations from non-AD, early-AD, and late-AD donors^5^ (Fig. 4A). Human population labels were retained from the source dataset. Cross-species comparisons were made between biologically related neuronal classes rather than by assuming one-to-one correspondence between mouse and human transcriptomic subtypes: mouse Ex1_IT was compared with human superficial IT and RORB-positive populations; Ex2_L5deep and Ex3_L6CT with deep-layer excitatory populations; Ex5_CLA_EP_Car3 with EX L5/6 IT Car3; In1_CGE_Calb2 with VIP/CGE-related populations; and In2_MGE_CGE_mix with broader SST-, PVALB-, and VIP-related interneuron classes. Because In2_MGE_CGE_mix contains mixed interneuron features at the resolution analyzed here, comparisons involving this population were made at the class rather than subtype level. LAMP5 and Rosehip-related populations were included as prespecified human comparators for testing the neuronal selectivity of mouse-derived program engagement.

**Figure 4.**
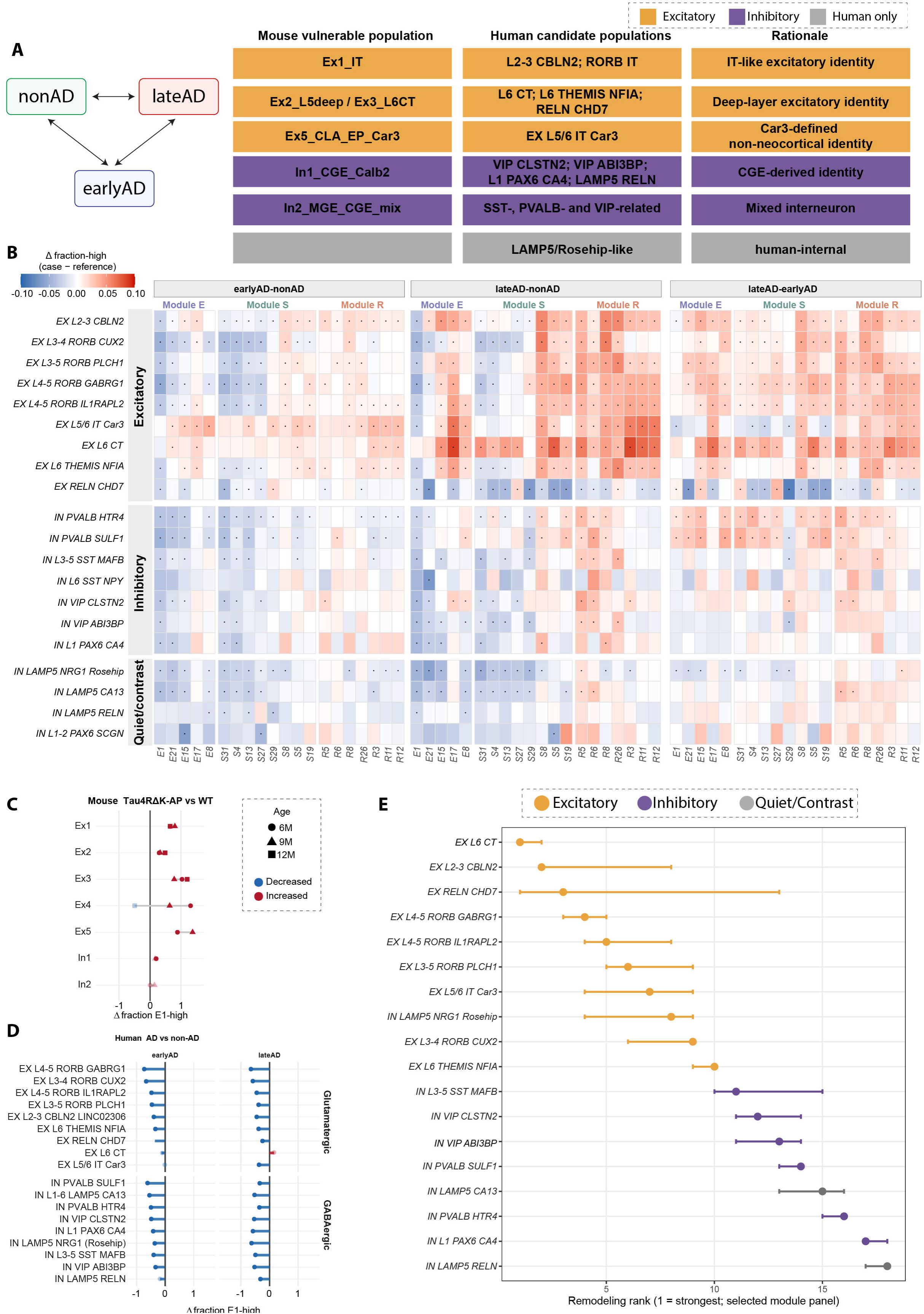
Mouse-derived neuronal-state programs show selective and stage-dependent engagement in human Alzheimer’s disease neurons. **(A)** Cross-species framework for evaluating mouse-derived neuronal-state programs in human AD. Human populations retain published source-dataset nomenclature and were related to mouse populations at the level of broad neuronal identity rather than by assuming one-to-one transcriptomic subtype equivalence. LAMP5 and Rosehip-related populations were included as human comparator identities. **(B)** Disease-stage-associated remodeling of mouse-derived module states across human neuronal populations. Heatmaps show case-minus-reference differences in donor-level module-high occupancy for earlyAD versus nonAD, lateAD versus nonAD, and lateAD versus earlyAD. Rows denote annotated human neuronal populations and columns denote selected E-, S-, and R-family modules. Red indicates higher and blue lower module-high occupancy in the case group. Black dots denote donor-level FDR < 0.05. Human donor, rather than nucleus, was the unit of biological inference. **(C)** E1 remodeling across mouse Tau4RΔK-AP neuronal populations. Points show the Tau4RΔK-AP-minus-WT difference in E1-high fraction for the indicated neuronal identities and ages. Point shape denotes age and color denotes direction of effect. E1 expansion is concentrated in glutamatergic identities at the well-sampled 6- and 9-month comparisons, while Ex4_DG_like reverses direction by 12 months. **(D)** E1 remodeling across selected human neuronal populations in early and late AD relative to nonAD. Points show the donor-level difference in E1-high occupancy. Negative values indicate reduced E1 engagement in AD. Populations are grouped by glutamatergic and GABAergic identity. **(E)** Ranking of human neuronal populations by aggregate lateAD-associated remodeling across the selected module panel. Rank order is determined by the unweighted mean absolute donor-level lateAD-versus-nonAD difference in module-high occupancy, with rank 1 indicating the strongest aggregate remodeling. Point color denotes neuronal class. Horizontal ranges show the range of ranks obtained by leave-one-donor-out sensitivity analysis.

Human AD neuronal populations showed pronounced selectivity in remodeling of the mouse-derived programs (Fig. 4B; Table S12). Changes were broadest in several excitatory identities, including EX L6 CT, EX RELN CHD7, EX L2-3 CBLN2, and multiple RORB-positive populations. Inhibitory populations generally showed narrower remodeling that depended on program family and stage-group comparison. Thus, engagement of the mouse-derived framework in human AD was structured by neuronal identity rather than being a uniform consequence of diagnosis.

### Human AD engages related neuronal-state programs with stage- and population-specific direction

Human AD did not reproduce the direction of every mouse-derived module effect (Fig. 4C, D; Fig. S8; Table S13). In Tau4RΔK-AP mice, E1 was among the most recurrent excitability-associated states and showed a strong glutamatergic bias at the better-sampled 6- and 9-month comparisons. All five glutamatergic populations included in the broader E1 analysis shifted positively at these ages, whereas the two inhibitory populations remained comparatively close to WT.

The direction of E1 was nevertheless not fixed within the mouse dataset. Ex4_DG_like showed strong E1 expansion at 6 months, remained increased at 9 months, and shifted below WT by 12 months. Thus, module direction was not fixed across disease stage within the same neuronal identity.

Human AD showed a predominantly opposite E1 pattern. Across the selected populations, E1-high occupancy was generally lower in early AD relative to non-AD and remained predominantly reduced in late AD, with only limited positive exceptions (Fig. 4D; Fig. S8). This decrease extended across excitatory and inhibitory classes and did not reproduce the glutamatergic-selective pattern observed in Tau4RΔK-AP mice.

Other E-family programs showed different behavior. For example, EX L5/6 IT Car3 remained near zero to negative for E1 while increasing in several other excitability-associated programs in early AD. Thus, cross-species correspondence at the level of broader biological program families did not require concordant direction of individual modules.

### Late-AD groups show increased engagement of regulatory and alternative synaptic programs

The late-AD group showed a remodeling architecture distinct from both non-AD and early-AD groups rather than simply stronger expression of the E-family states that were reduced earlier (Fig. 4B). Several excitatory populations showed increased engagement of R-family programs in late AD relative to non-AD and in late-AD relative to early-AD comparisons. Broad regulatory remodeling was evident in EX L6 CT, EX L5/6 IT Car3, EX RELN CHD7, and multiple RORB-positive populations, involving programs including R5, R6, R8, R26, R3, R11, and R12. Selected synaptic and excitability-associated programs were also increased in these populations.

Inhibitory populations showed a more restricted stage-associated pattern. PVALB-, SST-, VIP/CGE-, and L1 PAX6-related populations generally showed decreases in selected E- and S-family states in early AD relative to non-AD, followed by more selective regulatory engagement in late AD, particularly among R5-, R6-, and R26-associated programs (Fig. 4B).

### State remodeling is concentrated in human excitatory populations independently associated with vulnerability

We next asked whether the human populations showing the strongest engagement of the mouse-derived programs corresponded to neuronal identities independently implicated in AD vulnerability (Fig. 4E). Eighteen populations were ranked using the mean absolute donor-level late-AD versus non-AD occupancy effect across the selected module panel, with leave-one-donor-out analysis used to assess rank stability.

All four RORB-positive excitatory populations ranked within the top ten. RORB-positive excitatory neurons have independently been implicated in selective vulnerability in human AD^7^. Thus, the strongest human engagement of the mouse-derived state architecture was concentrated in neuronal identities associated independently with vulnerability, even though neither RORB status nor depletion was used to define the modules.

The prespecified LAMP5 comparators showed a different pattern (Fig. S9). IN LAMP5 CA13 and IN LAMP5 RELN occupied the low-remodeling end of the ranking, whereas the human-specific IN LAMP5 NRG1 Rosehip population showed intermediate remodeling dominated by negative effects. Closely related inhibitory populations therefore differed substantially in their engagement of the mouse-derived programs.

### A depletion-associated human identity program overlaps selected synaptic and excitability-associated states

The modules analyzed here were nominated from disease-associated transcriptional changes and describe states that change with pathology. We next asked whether the same framework also intersected constitutive molecular features of a neuronal identity independently associated with depletion in human AD. We compared the fixed module panel with the control-derived Exc NRGN BEX1 identity program reported by Pereira et al.^8^ as reproducibly depleted across clinical AD presentations (Fig. S10; Table S14).

The overlap extended across synaptic, postsynaptic, and selected calcium-signaling components of the framework. S18 and S11 were among the strongest synaptic enrichments, while the nested NMDAR-associated modules S13 and S2 and the receptor-gating module S19 were also enriched. Several E-family programs, including E15 and E13, showed significant overlap. Recurrently shared genes included CAMK2A, CAMK2B, NRGN, NEFL, PRKACB, SNAP25, VAMP2, STXBP1, NSF, NPTN, and TSPAN7.

E1 itself was not significantly enriched, and R13 showed no corresponding overlap. The depletion-associated human identity therefore intersected the broader synaptic and calcium-signaling architecture remodeled in disease without containing every disease-associated state. This separation supports a distinction between constitutive molecular features associated with depletion and transcriptional states superimposed on neuronal identity during disease.

Taken together, mouse-derived neuronal-state programs were engaged in human AD in a population-selective and stage-associated manner, with prominent remodeling across synaptic, excitability-associated, and regulatory programs in several excitatory identities independently associated with vulnerability (Fig. S10C, D). Cross-species correspondence was strongest at the level of broader biological architecture rather than the direction of every individual module.

Independently defined human depletion-associated identity mapped onto selected synaptic, receptor-proximal, and calcium-signaling components of this architecture, while excluding other disease-responsive modules. Thus, constitutive vulnerability-associated identity and disease-associated neuronal state intersect, but are not equivalent.

### An independent tau model shows related population-specific state remodeling across cortex and hypothalamus

To test whether the state architecture recurred across a genetically distinct amyloid-tau model and RNA-capture modality, we profiled cortex and hypothalamus from 6-month-old male WT and TauP301S-AP (TauP301S crossed with APP;PS1) mice by single-nucleus RNA sequencing (Fig. 5A, referred to as WT and TauAPP). This dataset differed from the discovery cohort in tau construct, RNA-capture modality, sex, regional sampling, and clustering resolution and was therefore analyzed separately rather than pooled. Neuronal populations were annotated from marker expression and broad biological identity without assuming one-to-one correspondence with populations in the Tau4RΔK-AP discovery cohort (Fig. 5B; Fig. S11, Tables S15, 16).

**Figure 5.**
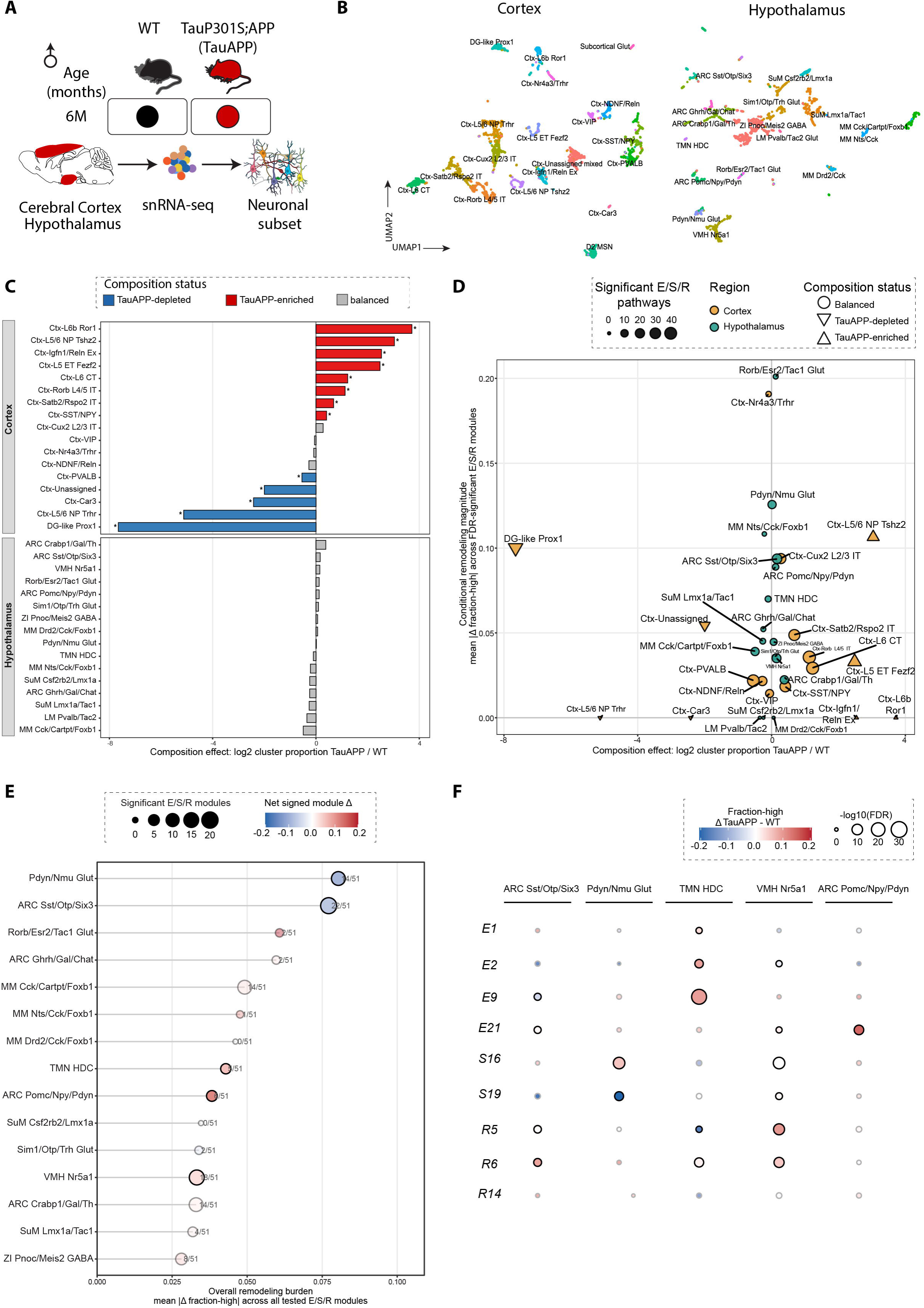
An independent tau model reproduces state remodeling and extends the framework beyond cortex. **(A)** Experimental design for the independent TauP301S-AP cohort. Cortex and hypothalamus from 6-month-old male WT and TauP301S-AP mice were profiled by single-nucleus RNA sequencing. **(B)** UMAP representations of cortical and hypothalamic neuronal nuclei annotated by marker expression, reference-assisted annotation, and broad biological identity. **(C)** Changes in relative neuronal representation in cortex and hypothalamus. Bars show the log2 fold change in population representation in TauP301S-AP relative to WT. Populations with log2FC ≤ -0.5 are shown as decreased, populations with log2FC ≥ 0.5 as increased, and populations between these boundaries as comparatively balanced. **(D)** Relationship between relative neuronal representation and transcriptional-state remodeling across cortical and hypothalamic populations. The x-axis shows population-representation log2 fold change and the y-axis shows mean absolute Δfraction-high across the tested E-, S-, and R-family modules. Point color denotes brain region, point shape denotes descriptive representation category, and point size denotes remodeling breadth, defined as the number of tested modules with |Δfraction-high| ≥ 0.05. **(E)** Transcriptional-state remodeling across hypothalamic neuronal populations. The x-axis shows mean absolute Δfraction-high across tested E-, S-, and R-family modules. Point size denotes remodeling breadth, defined as the number of modules with |Δfraction-high| ≥ 0.05. Point color summarizes the mean signed Δfraction-high among modules meeting this effect-size threshold, with positive and negative values indicating remodeling dominated by increased or reduced module-high occupancy, respectively. **(F)** Selected module-state effects across hypothalamic neuronal populations, including ARC Sst/Otp/Six3, Pdyn/Nmu Glut, TMN HDC, VMH Nr5a1, and ARC Pomc/Npy/Pdyn. Rows show selected E-, S-, and R-family modules. Circle color represents signed TauP301S-AP-minus-WT Δfraction-high and circle size represents absolute effect magnitude, |Δfraction-high|.

Population correspondence between cohorts was asymmetric in resolution. The discovery Ex1_IT population encompassed IT-like identities resolved here as Ctx-Cux2 L2/3 IT, Ctx-Rorb L4/5 IT, and Ctx-Satb2/Rspo2 IT. Ctx-L6 CT (Foxp2/Tle4/Syt6) provided a deep-layer corticothalamic counterpart to Ex3_L6CT. The deep-layer compartment was resolved further in this cohort than in the discovery data: Ctx-L5 ET Fezf2 represents a corticofugal-like identity, while Ctx-L5/6 NP Trhr and Ctx-L5/6 NP Tshz2 represent two distinct near-projecting-like glutamatergic populations. No single population in this cohort corresponds directly to the discovery Ex2_L5deep annotation, and we therefore treat these deep-layer identities as related at the class level only. DG-like Prox1 corresponded broadly to Ex4_DG_like, and Ctx-Car3 to the Ex5_CLA_EP_Car3 class, although transcriptomic identity alone does not establish a claustrum or endopiriform anatomical origin. Interneuron identities were resolved more finely into Ctx-SST/NPY, Ctx-PVALB, Ctx-NDNF/Reln, and Ctx-VIP populations, so cross-cohort inhibitory comparisons were made at the class rather than subtype level.

### Cortical relative representation does not predict remodeling magnitude

Differences in cortical relative representation occurred in both directions across TauP301S-AP neuronal populations (Fig. 5C). Large positive shifts were observed for Ctx-L6b Ror1, Ctx-L5/6 NP Tshz2, Ctx-Igfn1/Reln Ex, and Ctx-L5 ET Fezf2, with more moderate positive shifts in Ctx-L6 CT, Ctx-Rorb L4/5 IT, Ctx-Satb2/Rspo2 IT, and Ctx-SST/NPY. DG-like Prox1 and Ctx-L5deep Trhr showed the largest negative shifts, while Ctx-Car3 and Ctx-PVALB were also reduced. Ctx-Cux2 L2/3 IT, Ctx-VIP, Ctx-NDNF/Reln, and Ctx-Nr4a3/Trhr remained comparatively close to WT.

Joint consideration of relative representation and transcriptional-state remodeling reproduced the broader non-redundancy observed in the discovery cohort (Fig. 5D; Table S17). Ctx-Nr4a3/Trhr responsive showed the strongest cortical remodeling despite remaining close to WT in relative representation. Ctx-L5deep Trhr showed the opposite configuration, with a large negative representation shift but little detectable remodeling. DG-like Prox1 also showed a large negative representation shift but, unlike Ctx-L5deep Trhr, underwent substantial molecular remodeling. Thus, populations with similarly reduced representation could differ markedly in the extent of within-population state change.

A similar contrast was evident among populations with positive representation shifts. Ctx-L5/6 NP Tshz2 combined a large positive shift in relative representation with strong remodeling, whereas Ctx-L6b Ror1 showed a comparably large positive representation shift but comparatively little remodeling. Across cortical populations, relative representation therefore provided limited information about the magnitude of transcriptional-state remodeling within the same neuronal identities.

Ctx-Nr4a3/Trhr responsiveness is partly defined by activity-associated genes, raising the possibility that immediate-early transcription contributes to its distinctive profile. Genotype-associated immediate-early gene expression across populations was therefore examined separately (Fig. S12).

### Related biological programs recur across models without fixed module-level direction

Focused module analysis showed that excitability-associated, synaptic, and regulatory remodeling recurred in the independent cortical cohort, although the particular programs and their directions differed among neuronal identities (Fig. S12A).

Ctx-Nr4a3/Trhr responsive, the cortical population with the strongest aggregate remodeling, showed coordinated changes across excitability-associated and regulatory programs, including E1, E15, and R14. Ctx-Rorb L4/5 IT showed a distinct profile dominated by E9 together with additional synaptic and excitability-associated effects, while Ctx-Cux2 L2/3 IT showed prominent E18 remodeling and changes across both E- and S-family programs. Related glutamatergic populations therefore engaged different components of the broader state architecture.

Ctx-L6 CT provided a particularly informative cross-model comparison. Its remodeling included decreases in E1, E18, E20, and S6 together with increases in E17 and several regulatory programs, including R11, R12, and R14. The related Ex3_L6CT population in the Tau4RΔK-AP discovery cohort instead showed positive E1 remodeling. Thus, the two models did not reproduce the direction of E1 within corticothalamic-like neurons, even though both showed substantial remodeling across related excitability-associated and regulatory programs.

Across the cortical dataset, several of the largest effects involved modules that were not dominant in the discovery cohort (Fig. S12B, C). The independent model therefore reproduced the broader organization of population-specific remodeling rather than a fixed module signature.

### Hypothalamic populations show substantial state remodeling despite comparatively preserved representation

Hypothalamic populations remained comparatively close to WT in relative representation, without the large bidirectional composition shifts observed in cortex (Fig. 5C, Table S17). Despite this relative compositional stability, several hypothalamic identities exhibited substantial remodeling of molecular states (Fig. 5D, E). The hypothalamus therefore provided a regional extension of the representation-state dissociation in populations whose relative representation was largely preserved.

Remodeling breadth varied markedly across hypothalamic identities. Pdyn/Nmu Glut and ARC Sst/Otp/Six3 showed the broadest responses across the tested module panel, while Rorb/Esr2/Tac1 Glut also showed strong conditional remodeling. Other populations, including VMH Nr5a1, ARC Crabp1/Gal/Th, ARC Ghrh/Gal/Chat, and TMN HDC, showed more intermediate or focused responses. Remodeling breadth and net direction were not equivalent, indicating that populations could differ both in how many programs were altered and in whether those changes were predominantly positive or negative.

Selected module effects further demonstrated population-specific organization (Fig. 5F). ARC Sst/Otp/Six3 showed changes across excitability, synaptic, and regulatory programs, including reduced E2 and S19 together with increased R6. Pdyn/Nmu Glut was distinguished by opposing synaptic effects, with increased S16 and reduced S19. VMH Nr5a1 showed prominent regulatory remodeling, while ARC Pomc/Npy/Pdyn showed a more focused response that included increased E21.

The newly resolved hypothalamic annotations also identified ARC Ghrh/Gal/Chat as a Ghrh/Gal/Chat GABA/cholinergic-like population and LM Pvalb/Tac2 Glut as a Pvalb/Tac2/Grin2c lateral-mammillary-like glutamatergic population. These identities further illustrate the cellular diversity captured by the hypothalamic dataset but were not the principal populations driving the remodeling patterns described above.

### TMN HDC neurons show focused excitability-associated remodeling despite preserved relative representation

TMN HDC neurons, corresponding to Hdc-positive tuberomammillary histaminergic neurons, showed a comparatively focused remodeling profile rather than the broad response observed in Pdyn/Nmu Glut or ARC Sst/Otp/Six3 (Fig. 5E, F). Their strongest selected effect was an increase in E9, accompanied by increased E2 and a smaller positive E1 shift. Selected synaptic and regulatory programs moved in the opposite direction, including S16 and R5. TMN HDC neurons combined focused excitability-associated remodeling with comparatively preserved relative representation.

This profile extends the state-remodeling framework to a neuromodulatory hypothalamic population without requiring the same module combination observed in cortex. Together with the broader hypothalamic results, it shows that substantial within-identity remodeling can occur in a region where relative representation remains comparatively close to WT.

Overall, the TauP301S-AP dataset reproduced the broader relationship between relative representation and within-population molecular state under an independent experimental configuration that differed in tau construct, sex, RNA-capture modality, regional sampling, and clustering resolution. In cortex, populations with similar representation changes could show markedly different degrees of remodeling, while strong remodeling also occurred in populations whose representation remained close to WT. The hypothalamic analysis extended this framework beyond cortex, with comparatively stable relative representation but substantial remodeling in selected neuronal identities. TMN HDC neurons provided a clear example of focused excitability-associated remodeling in a population with comparatively preserved representation. Together, these findings show that the separation between population representation and molecular state recurs across neuronal identities and brain regions, while the specific transcriptional programs involved remain context-dependent.

### AMPK activation under oxidative stress shifts the E1-associated transcriptional state

To test whether a disease-associated transcriptional state identified in vivo could be experimentally perturbed in a neuronal-like cellular context, we used retinoic acid-conditioned Neuro2a cells exposed to oxidative stress and selected signaling perturbations (Fig. 6A). Retinoic acid conditioning produced broad transcriptional changes associated with membrane regulation, cellular signaling, and neuronal differentiation (Fig. S13A, C). smFISH independently confirmed MAP2 and TUBB3 expression, including neurite-like processes (Fig. S13B). Retinoic acid also increased aggregate expression of selected synaptic and excitability-associated module families (Fig. S13D), consistent with acquisition of a partial neuronal-like state.

**Figure 6.**
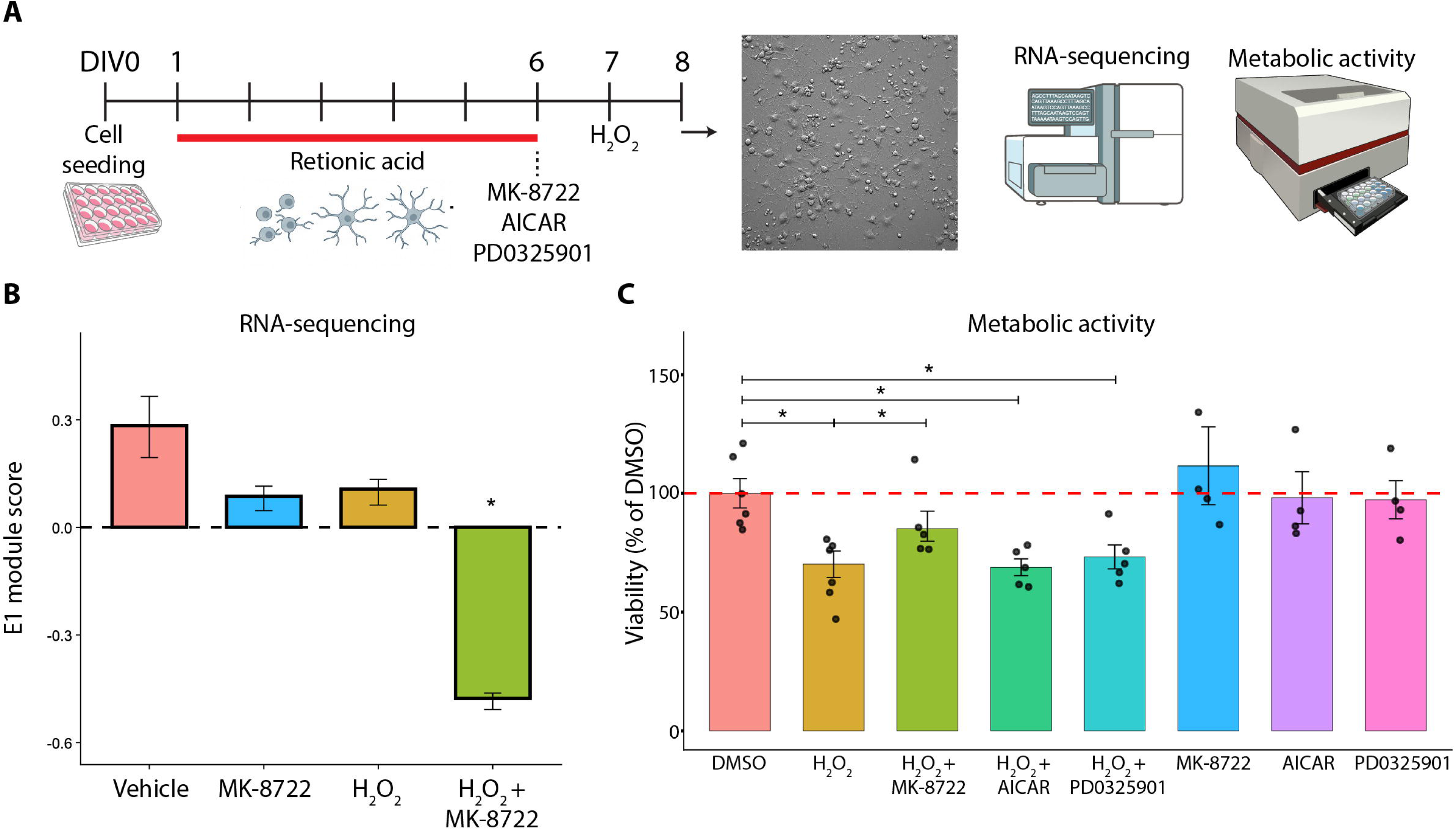
Direct AMPK activation reduces the E1-associated transcriptional program under oxidative stress in retinoic acid-conditioned Neuro2a cells. **(A)** Experimental design. Neuro2a cells were seeded at 2.0 × 10 cells per well on poly-D-lysine-coated 24-well plates (DIV0). From DIV1, cells were conditioned in DMEM containing 2% FBS and 30 µM all-trans retinoic acid (0.03% final DMSO), with medium replaced every other day. On DIV6, cells were pretreated for 2 h with 400 nM MK-8722, 40 µM AICAR, 400 nM PD0325901, or matched vehicle, followed by addition of 500 µM H O or vehicle without medium replacement and a further 24 h incubation. Cells were then processed for phase-contrast imaging, bulk RNA-sequencing, and PrestoBlue metabolic-activity assay. **(B)** Sample-level module score for the fixed E1 gene set (NMDAR-linked AMPK signaling program) in the four-condition bulk RNA-sequencing design (vehicle, MK-8722, H O, H O + MK-8722). Scores are the mean gene-wise standardized log (CPM + 1) across represented E1 genes. Bars show mean ± SEM. *p < 0.05. This score is a distinct quantity from the single-cell AUCell/module-high fraction (ΔF) with the same fixed gene set. **(C)** Metabolic activity measured by PrestoBlue HS after 24 h of H O exposure, background-corrected against cell-free wells and expressed relative to the mean vehicle (DMSO) signal of the same experiment (red dashed line, 100%). Bars show mean ± SEM. Horizontal bars indicate comparisons with DMSO; *p < 0.05.

H O exposure induced a strong oxidative-stress transcriptional response, including induction of *Ddit3*, *Hmox1*, and *Atf3* (Fig. S13E). Addition of the direct AMPK activator MK-8722 under H O exposure produced a distinct transcriptional response (Fig. S13F, G), including decreased expression of several neuronal-associated transcripts such as *Dlg4*, *Camk2a*, *Stmn2*, *Snap25*, *Gap43*, *Map2*, and *Tubb3*.

We next quantified the fixed E1 gene set in the bulk RNA-seq samples using the same finalized E1 definition applied in the single-cell analyses (Fig. 6B; Table S18). The combined H O and MK-8722 condition showed a marked reduction in the E1-associated transcriptional score relative to the other RNA-seq conditions, whereas MK-8722 alone and H O alone did not produce the same shift. Thus, direct AMPK activation in the context of oxidative stress shifted the E1-associated transcriptional state in the negative direction.

Metabolic activity was assessed independently using PrestoBlue (Fig. 6C). H O reduced metabolic activity relative to DMSO, whereas MK-8722 partially restored it under oxidative stress. AICAR and the MEK inhibitor PD0325901 were included as comparator perturbations in this assay and did not produce a comparable restoration under the conditions tested. MK-8722 alone did not reduce metabolic activity. The transcriptional and metabolic responses therefore diverged under combined H O and MK-8722 treatment, with reduced E1-associated transcription accompanied by partial recovery of metabolic activity.

Together, these experiments show that the E1-associated transcriptional state is experimentally modifiable and that its direction depends on signaling context rather than tracking AMPK activation in a simple monotonic manner.

## Discussion

### Neuronal involvement is not captured by relative representation alone

Selective neuronal vulnerability is commonly inferred from which populations are depleted in disease. Our data indicate that this captures only one dimension of neuronal involvement. Across Tau4RΔK-AP cortex, substantial transcriptional remodeling occurred in populations whose relative representation remained close to WT or increased. In contrast, some populations with large decreases in representation showed comparatively limited remodeling among the neurons recovered within them. Relative representation and molecular state are non-redundant measurements, although the present data do not establish that they are biologically independent.

This distinction is relevant because maintenance of neuronal identity and function depends on active transcriptional and chromatin-based regulation^24^. Disruption of these systems can produce molecular and functional abnormalities before, or in the absence of, overt neuronal loss^11,12^, and functional impairment can occur despite preserved structural representation^9^. Consistent with this distinction, early transcriptional remodeling across the five populations available for age-structured analysis did not provide a robust monotonic ordering of representation differences at later ages. We therefore view relative representation, the magnitude of transcriptional remodeling, and the distributional form of that remodeling as complementary features of neuronal involvement rather than successive stages of a demonstrated trajectory.

### Synaptic and excitability programs define recurrent but context-dependent axes of remodeling

Synaptic remodeling was not characterized by a uniform decline in synaptic gene expression. Instead, the NMDAR-associated hierarchy showed a positional reorganization in which downstream postsynaptic signaling programs increased while receptor-proximal gating and trans-synaptic adhesion programs decreased. Because several of these Reactome modules are hierarchically nested, this pattern is better interpreted as coordinated reorganization of a shared signaling architecture than as independent pathway effects. Such organization is consistent with broader evidence implicating synaptic function and plasticity-associated genes in vulnerable human AD neuronal identities and in modulation of tau toxicity^8,10,19^.

Excitability and Ca² -associated programs formed another recurrent axis. E1 was among the prominent positive states in the Tau4RΔK-AP discovery cohort, particularly in glutamatergic populations at 6 and 9 months, and connects CAMKK2-AMPK signaling with downstream microtubule-associated biology. The underlying pathway is relevant to AD, as CAMKK2-AMPK signaling has been linked experimentally to Aβ-associated tau phosphorylation and synaptic dysfunction ^22,23^, while neuronal hyperactivity and Ca² dysregulation are established features of experimental amyloid pathology^25,26^.

The direction of individual states, however, was not fixed. E1 increased recurrently in Tau4RΔK-AP glutamatergic neurons but decreased in TauP301S-AP corticothalamic-like neurons, was predominantly reduced in human AD, and reversed direction with age in Ex4_DG_like. The recurrent feature across datasets was therefore engagement of related excitability and Ca² -associated biology rather than conservation of a single module or its sign. More broadly, these findings argue against treating any individual transcriptional module as a universal marker of neuronal vulnerability or biochemical pathway activity.

### Disease changes the distribution of neuronal states, not only their average

Population averages and threshold occupancy did not fully capture the forms of state remodeling observed at single-cell resolution. Disease displaced cellular-state distributions in some comparisons, broadened them in others, and narrowed them in still others. These configurations imply different forms of within-identity remodeling: coordinated displacement reflects a population-wide shift in state, broadening indicates greater heterogeneity among neurons assigned to the same identity, and narrowing indicates convergence toward a more restricted range of states.

The late R13 response in In1_CGE_Calb2 illustrates narrowing, with a large increase in module-high occupancy accompanied by a comparatively smaller central displacement and reduced variance. By contrast, the Ex3_L6CT E1 distribution at 12 months shifted toward higher values while broadening, whereas In2_MGE_CGE_mix at 6 months showed modest E1 displacement with essentially unchanged q90 occupancy. Threshold occupancy, central displacement, and distributional width therefore captured partly separable features of remodeling. Together with the opposing movement of different programs within the same neuronal identities, these observations indicate that disease reorganizes neuronal state in a structured rather than uniformly destabilizing manner.

The chromatin analysis captured selected components of this organization. Accessibility changes were most directionally concordant with RNA for the declining receptor-gating and adhesion programs, whereas increasing downstream postsynaptic modules and E1 lacked a uniform GeneActivity counterpart. RNA and ATAC effects were also not globally associated across matched comparisons. Cross-modal correspondence was therefore selective, suggesting that some transcriptional changes are accompanied by detectable accessibility remodeling while others are not proportionally reflected by GeneActivity.

### An independent cohort recapitulates the separation between representation and molecular state

The independent TauP301S-AP cohort reproduced the broader organization of neuronal remodeling despite differences in tau construct, sex, RNA-capture modality, regional sampling, and clustering resolution. Cortical glutamatergic populations again showed remodeling across excitability, synaptic, and regulatory programs, while the magnitude of molecular-state change remained incompletely coupled to relative representation.

This separation was evident across populations with contrasting outcomes. Ctx-L5deep Trhr and DG-like Prox1 both decreased substantially in representation but differed markedly in molecular remodeling, whereas Ctx-Nr4a3/Trhr responsive showed strong state remodeling while remaining comparatively close to WT representation. Related populations also differed in the particular modules and directions involved, including opposite E1 direction in corticothalamic-like populations across the two models. Thus, the independent cohort reproduced the broader relationship between representation and state without requiring a fixed transcriptional signature.

Experimental and biological differences may contribute to module-level discrepancies between cohorts, including differences in whole-cell versus nuclear RNA recovery, tau construct, sex, stage, dissection, and clustering resolution^17,18^. These factors cannot be separated individually here. Importantly, however, the broader separation between population representation and within-identity remodeling recurred under substantially different experimental conditions.

### Human Alzheimer’s disease engages related neuronal populations and biological programs

Projection of the fixed mouse-derived modules into human AD identified selective rather than global engagement. Remodeling was strongest in several deep-layer and RORB-positive excitatory populations independently implicated in human AD vulnerability, whereas inhibitory populations generally showed narrower and subtype-dependent changes. The prespecified LAMP5 comparison further illustrated this specificity: IN LAMP5 CA13 and IN LAMP5 RELN occupied the low-remodeling end of the ranking, whereas the human-specific IN LAMP5 NRG1 Rosehip population showed greater, predominantly negative remodeling. Closely related inhibitory identities were therefore not interchangeable in their engagement of the mouse-derived programs.

Comparison with an independently defined depletion-associated human identity program further separated constitutive identity features from disease-associated state remodeling. The Exc NRGN BEX1 program^8^, defined from control tissue independently of the present disease-state analysis, was enriched for synaptic, postsynaptic, NMDAR-proximal, and selected calcium/excitability-associated modules, including S18, S11, S13, S2, and S19 together with several E-family programs. Other disease-responsive modules, including E1 and R13, were not similarly enriched. The depletion-associated identity therefore intersected selected components of the broader disease-remodeled architecture without reproducing it in full, supporting a distinction between constitutive vulnerability-associated identity and states superimposed during disease.

Individual module direction was less conserved across species. E1, for example, increased predominantly in Tau4RΔK-AP glutamatergic neurons but was generally reduced in human AD, with directional divergence also evident across mouse model and age. More broadly, related synaptic, excitability/Ca², and regulatory programs were repeatedly engaged, and the strongest human remodeling occurred in several neuronal identities independently associated with vulnerability. Cross-species correspondence therefore lay more strongly in the neuronal populations and biological programs involved than in a conserved direction or combination of every module.

Differences in disease duration, aging, survivorship, cellular context, and RNA-capture modality could all contribute to directional divergence between experimental models and human postmortem tissue. Age-associated epigenomic remodeling may further alter the regulatory landscape on which pathology acts^27^. The available data do not resolve these contributions, but they argue against a simple correspondence in which a single early module state determines later representation change or retains the same direction across disease contexts.

### State remodeling extends beyond cortex

The hypothalamic analysis extended this framework to a region in which relative neuronal representation remained comparatively stable. Several hypothalamic identities nevertheless showed substantial transcriptional remodeling, demonstrating that strong state changes are not restricted to regions with prominent compositional shifts. Hdc-positive TMN histaminergic neurons illustrated a focused form of this response, with selected excitability and Ca² -associated programs changing despite comparatively preserved representation. Because TMN histaminergic neurons participate in arousal and wake regulation, these findings identify a potentially relevant neuromodulatory population in the context of processes disrupted in AD and other neurodegenerative disorders^28–30^, although the functional consequences of the molecular changes remain to be established.

### The E1-associated transcriptional state is pharmacologically modifiable under oxidative stress

The Neuro2a perturbation experiments provided an experimental extension of the *in vivo* state analysis. In retinoic acid-conditioned cells, direct AMPK activation with MK-8722 under oxidative stress reduced the E1-associated transcriptional score rather than reproducing the positive E1 shifts observed in Tau4RΔK-AP excitatory neurons. This change occurred alongside partial recovery of metabolic activity, indicating that the transcriptional and metabolic readouts did not move in parallel.

E1 represents transcription of genes within a curated Reactome pathway rather than a biochemical measurement of AMPK activity. The perturbation therefore demonstrates that the E1-associated transcriptional state is pharmacologically modifiable and that its direction does not map monotonically onto direct AMPK activation in this cellular context. Its variable direction across perturbation, age, model, and human disease further emphasizes that pathway-associated transcriptional states are context dependent rather than fixed indicators of pathway activity. One possibility is that sustained pathway engagement elicits transcriptional feedback that changes expression of components represented in the module, a model that could be tested directly by measuring E1 transcription together with AMPK and ACC phosphorylation across a perturbation time course.

### Scope and limitations

Several features of the study constrain interpretation. First, dissociated-cell and isolated-nucleus datasets measure relative recovery rather than absolute neuronal number. The pronounced reduction in recovered inhibitory neurons in 12-month Tau4RΔK-AP cortex therefore requires independent histological validation before being interpreted as neuronal loss, and sparse inhibitory recovery at this age limits state-level analysis, particularly for In2_MGE_CGE_mix.

Second, the progression design is cross-sectional at the level of individual neurons. It permits comparison of population-level states across ages but cannot determine the fate of neurons occupying particular remodeled states. The age-structured analysis included only five populations and excluded a strong monotonic relationship between early remodeling and later representation change rather than establishing temporal independence.

Third, the S, E, and R families are organizational groupings of overlapping and sometimes hierarchically related Reactome pathways rather than statistically independent biological systems. Gene sharing and pathway nesting must therefore be considered when interpreting coordinated effects.

Fourth, the primary module panel was nominated from differential expression in the discovery cohort. Complete curated Reactome gene sets were subsequently fixed and applied unchanged across downstream analyses, including the independent mouse cohort and human datasets. This limits circularity in subsequent testing but retains the discovery bias introduced by pathway nomination, and biological processes outside the nominated panel would not be captured.

Fifth, cross-dataset comparisons are influenced by differences in tau construct, sex, pathological stage, brain dissection, clustering resolution, and RNA-capture modality. The primary Tau4RΔK-AP progression cohort was female, whereas the complementary 12-month Tau4RΔK-AP cohort and independent TauP301S-AP cohort were male. The study was not designed to resolve sex-specific trajectories from cohort or model effects.

Finally, the human analyses are cross-sectional and shaped by donor heterogeneity, aging, disease duration, and survivorship. They establish engagement of related biological programs in human AD but cannot determine whether differences between diagnostic groups represent transitions experienced by individual neurons.

## Conclusion

Relative neuronal representation and molecular state capture distinct dimensions of disease-associated neuronal involvement. Across amyloid-tau models and human AD, neuronal identities showed structured, population-specific remodeling of synaptic, excitability-associated, and regulatory programs that did not consistently track changes in relative representation. Thus, selective neuronal involvement is defined not only by which populations remain represented in disease, but also by the molecular states occupied by those neurons.

## Methods

### 1. Animals and tissue collection

#### 1.1 Tau4RΔK-AP disease-progression cohort

Previously generated single-cell RNA-sequencing (scRNA-seq) and single-cell chromatin-accessibility (scATAC-seq) datasets from the Tau4RΔK-AP disease-progression cohort were reanalyzed in the present study^20^. The cohort comprised four genotypes: wild-type (WT), APP;PS1, Tau4RΔK, and Tau4RΔK-AP. Tau4RΔK-AP mice carried the APPswe, PS1ΔE9, CamKII-tTA, and TetO-TauRDΔK transgenes; Tau4RΔK mice carried CamKII-tTA and TetO-TauRDΔK; and APP;PS1 mice carried APPswe and PS1ΔE9.

Tau4RΔK mice were generated by crossing TetO-TauRDΔK mice, which express a mutant tau repeat-domain construct under tetracycline-responsive control, with CamKII-tTA driver mice. Tau4RΔK mice were subsequently crossed with APPswe;PS1ΔE9 mice to generate Tau4RΔK-AP animals carrying both amyloid- and tau-associated transgenes.

Transgenic-line generation, genetic background, breeding, genotyping, husbandry, doxycycline regulation, and the original tissue-collection procedures have been described previously^20^.

The primary disease-progression analysis used cortical datasets from female mice collected at 6, 9, and 12 months of age across all four genotypes. Female mice were used for the progression series because the Tau4RΔK-AP phenotype was previously shown to be more pronounced in females^31^. A previously generated 12-month male cohort containing the same four genotypes was analyzed as a complementary biological cohort and was not used to infer a longitudinal or sex-specific disease trajectory.

For the scATAC-seq analysis, the female cohort comprised 13 biological samples: at 6 months, two WT, two Tau4RΔK, two APP;PS1, and three Tau4RΔK-AP samples; and at 12 months, one sample from each genotype.

#### 1.2 Independent TauP301S-AP single-nucleus RNA-sequencing cohort

An independent single-nucleus RNA-sequencing (snRNA-seq) cohort was generated at Aarhus University using 6-month-old male WT and TauP301S-AP mice.

The tau component was derived from heterozygous Tau P301S mice [B6;C3-Tg(Prnp-MAPT*P301S)PS19Vle/J], which express human *MAPT* carrying the P301S mutation under control of the prion protein promoter. The amyloid component was derived from heterozygous APP/PS1 mice [B6;C3-Tg(APPswe,PSEN1dE9)85Dbo/Mmjax], carrying the APP Swedish mutation and PSEN1ΔE9. Mice carrying both transgenic components are referred to throughout this study as TauP301S-AP. This cohort therefore represents a genetically distinct combined amyloid-tau model from the Tau4RΔK-AP discovery cohort.

#### 1.3 Animal housing and experimental conduct

Animals in the independent TauP301S-AP cohort were housed at the Aarhus University Department of Biomedicine animal facility under a 12-h light/dark cycle with *ad libitum* access to standard chow and water. Mice were group-housed in individually ventilated cages with environmental enrichment according to institutional husbandry procedures.

Because experimental groups were defined by genotype, random allocation to genotype was not applicable. Tissue collection, nuclei isolation, library preparation, sequencing, and primary bioinformatic processing were performed using the same procedures across WT and TauP301S-AP samples within each brain region.

#### 1.4 Tissue collection and regional dissection

For the previously generated Tau4RΔK-AP cohort, cerebral cortical tissue was collected and processed as described previously^20^.

For the independent TauP301S-AP cohort, 6-month-old male mice were euthanized by cervical dislocation, and brains were rapidly removed and dissected on an ice-cold surface. Cerebral cortex and hypothalamus were collected as separate regional specimens from the same animals. Cortical tissue was dissected according to the procedure described previously^20^, with neocortex separated from underlying subcortical structures. Hypothalamic tissue was dissected according to previously described anatomical boundaries^32^: the optic chiasm defined the rostral boundary, the mammillary bodies the caudal boundary, the hypothalamic sulci the lateral boundaries, and the ventral level of the third ventricle the dorsal boundary. Immediately after dissection, tissue was snap-frozen on dry ice and stored at −80°C until nuclei isolation.

#### 1.5 Animal ethics

Animal procedures for the independent TauP301S-AP cohort were conducted at Aarhus University in accordance with institutional requirements and applicable Danish and European legislation governing the use of animals for scientific purposes. Experiments were performed under authorization 2023-15-0201-01605 from the Danish Animal Experiments Inspectorate. Ethical approvals and procedures for the previously generated Tau4RΔK-AP cohort were reported with the original study^20^.

### 2. Single-cell RNA sequencing of the Tau4R**Δ**K-AP cohort

#### 2.1 Preparation of single-cell suspensions

Single-cell RNA-sequencing data from the Tau4RΔK-AP cohort were generated previously and reanalyzed in the present study. Cerebral cortical tissue was dissociated using papain-based enzymatic digestion followed by mechanical trituration, as described previously^20,32^. Briefly, freshly dissected cortical tissue was collected in Hibernate-A medium supplemented with 2% B-27 and 0.5 mM GlutaMAX and enzymatically dissociated with papain. Tissue was then mechanically triturated, filtered to remove undissociated material and aggregates, and subjected to density-based debris removal. Cells were washed, filtered, and assessed for concentration and viability before library preparation. RNase inhibitor was included during the final preparation steps for samples processed for scRNA-seq. Detailed reagent concentrations and tissue-processing procedures are provided in the original studies^20,32^.

#### 2.2 scRNA-seq library preparation and sequencing

Single-cell RNA-seq libraries were generated previously using the 10x Genomics Chromium platform as described for the original Tau4RΔK-AP cohort^20^. No new single-cell libraries from this cohort were generated for the present study. Library preparation, sequencing, and initial processing procedures are described in the original publication.

#### 2.3 Neuronal dataset construction and nomenclature

Previously processed Seurat objects from the Tau4RΔK-AP cortical scRNA-seq dataset were used as the starting point for the present analyses. Metadata describing genotype, age, sex, biological-sample identity, and neuronal-cluster assignment were retained. Neuronal annotations were evaluated using reference-based annotation with MapMyCells^33^ together with established marker expression and were consolidated to the resolution required for the present analyses. Reference assignments were used to support broad biological identity rather than to impose one-to-one correspondence with external transcriptomic subtypes.

The principal neuronal populations used for composition and state analyses were Ex1_IT, an intratelencephalic Rorb-associated excitatory population; Ex2_L5deep, a deep-layer L5 excitatory population; Ex3_L6CT, a layer 6 corticothalamic population; Ex4_DG_like, a dentate granule-like excitatory population; In1_CGE_Calb2, a CGE-derived Calb2-positive interneuron population; and In2_MGE_CGE_mix, a mixed interneuron population containing MGE- and CGE-associated features at the resolution analyzed here.

Additional neuronal identities, including Ex5_CLA_EP_Car3, D1 and D2 striatal spiny projection neurons, and Cajal-Retzius cells, were annotated but excluded from the principal composition analyses. Ex5_CLA_EP_Car3 was retained in selected transcriptional-state analyses where cell recovery permitted. Because In2_MGE_CGE_mix was intentionally analyzed at combined resolution, no subtype-level SST inference was made from this population in the principal analyses. For figure display, the six principal populations are abbreviated as Ex1, Ex2, Ex3, Ex4, In1, and In2 where space is limited.

### 3. Neuronal representation analysis

#### 3.1 Broad excitatory and inhibitory neuronal representation

Cellular representation was quantified at the biological-sample level as the fraction of all quality-controlled cortical cells assigned to the neuronal class of interest. The principal excitatory class comprised Ex1_IT, Ex2_L5deep, Ex3_L6CT, and Ex4_DG_like, and the inhibitory class comprised In1_CGE_Calb2 and In2_MGE_CGE_mix. The total number of quality-controlled cortical cells from each biological sample was used as the denominator. Proportions were not renormalized within the neuronal compartment.

For the female Tau4RΔK-AP disease-progression cohort, representation was evaluated separately at 6, 9, and 12 months, with APP;PS1, Tau4RΔK, and Tau4RΔK-AP mice compared with age-matched WT controls. The complementary male analysis was restricted to the 12-month cohort. Because recovery from dissociated tissue can vary with neuronal identity and tissue condition, these measurements were interpreted as relative representation rather than absolute neuronal number.

#### 3.2 Neuronal population-level representation

Population-level representation was evaluated for Ex1_IT, Ex2_L5deep, Ex3_L6CT, Ex4_DG_like, In1_CGE_Calb2, and In2_MGE_CGE_mix. Relative representation was calculated for each biological sample as the number of recovered cells assigned to a neuronal population divided by the total number of quality-controlled cortical cells from the same sample. Sample-population combinations with no recovered cells were retained as zero counts.

Representation effects were displayed as log2 fold changes relative to the corresponding WT group. Mean sample-level representation within each genotype was used, with a half-cell pseudocount applied only to stabilize descriptive effect-size estimates for low-frequency populations. The pseudocount was not included in statistical testing. WT reference values were calculated independently within each age in the female progression cohort and within the 12-month male cohort.

#### 3.3 Statistical testing of representation differences

Genotype-associated differences in neuronal representation were tested using biological samples as the unit of replication. For each neuronal class or population, the number of recovered cells assigned to that category and the number of all remaining quality-controlled cortical cells from the same sample were analyzed using a quasibinomial generalized linear model with a logit link. The quasibinomial model was used to accommodate overdispersion in sample-level proportions.

Each disease genotype was compared separately with the corresponding age-matched WT group. Female comparisons were performed independently at 6, 9, and 12 months, and the male comparison was restricted to 12 months. Statistical testing was performed only when sufficient biological replication was available in both groups.

Benjamini-Hochberg correction was applied within the corresponding broad-class and neuronal-population analysis families. Nominal p values are shown where reported in the Results, while adjusted values were used to evaluate robustness to multiple testing.

#### 3.4 Descriptive composition categories used in integrated analyses

For integrated representation-state analyses, neuronal representation was calculated as described above and retained as a separate dimension from transcriptional-state remodeling. Populations with log2FC ≤ −0.5 were described as decreased in representation, those with log2FC ≥ 0.5 as increased in representation, and those with |log2FC| < 0.5 as comparatively stable. These thresholds were used only for descriptive organization and did not determine statistical significance.

### 4. Single-Cell Module Scoring and Transcriptional State Analysis

#### 4.1 Pathway-level gene-set generation

Pathway-level gene sets were generated using a two-stage selection and curation procedure. Disease-associated differentially expressed genes were first identified in the Tau4RΔK-AP cortical discovery cohort for each disease genotype relative to age-matched WT. Transgene-associated and genotype-defining genes, including *Mapt*, were excluded before Reactome enrichment and were not reintroduced into the finalized scoring modules. The highest-ranking pathways were selected for subsequent analysis.

Selected Reactome pathways were organized into three biological families: synaptic organization, receptor composition, adhesion, and neurotransmission (S family); excitability, Ca² signaling, ion handling, and activity-associated signaling (E family); and transcriptional regulation and stress-associated signaling (R family). Finalized scoring modules were derived from the curated Reactome pathway membership after identifier mapping, ortholog conversion, removal of duplicate or unmapped entries, and exclusion of prespecified transgene-associated or genotype-defining genes. Modules were not restricted to the differentially expressed genes that originally nominated the pathway and were not collapsed into family-level gene sets.

Molecular identifiers were extracted from the Reactome MoleculeName field. UniProt and Ensembl identifiers were mapped to HGNC gene symbols using biomaRt, entries already represented as gene symbols were retained, and duplicate, missing, and unmapped identifiers were removed. Human gene sets were converted to mouse orthologs using homologene. Where multiple mouse orthologs were returned, the first mapped mouse symbol returned by the procedure was retained. Modules containing fewer than 10 mapped mouse genes were excluded.

Stable E-, S-, and R-prefixed module identifiers were assigned using a fixed lookup table and retained across downstream analyses. Within each dataset, a module was scored only when at least 10 genes from its finalized definition were represented in the corresponding expression matrix. The finalized module definitions were otherwise held constant across genotype, age, neuronal population, mouse model, profiling platform, and cross-species analyses.

Because the primary panel was discovery-informed, the complete Reactome mouse pathway collection was additionally evaluated as a sensitivity analysis. Full-Reactome gene sets were processed using the same exclusions, minimum represented-gene criterion, AUCell scoring procedure, and WT-referenced module-high definitions. E1 was ranked within each neuronal population and age relative to the complete pathway collection and to pathways of comparable gene-set size.

#### 4.2 Per-cell AUCell scoring

Module activity was quantified at single-cell resolution using AUCell. For the Tau4RΔK-AP cortical dataset, log-normalized RNA-assay expression values were used to generate cell-specific gene-expression rankings with AUCell_buildRankings, followed by module scoring with AUCell_calcAUC.

Scores were generated jointly across the neuronal dataset, yielding a continuous AUCell score for each retained module in each cell. Only modules with at least 10 represented genes in the corresponding expression matrix were analyzed.

#### 4.3 Definition and summarization of module-high states

Module-high states were defined using WT-referenced AUCell percentiles. For the primary Tau4RΔK-AP analysis, the threshold for each module was set at the 90th percentile of the WT AUCell distribution within the corresponding neuronal population and age and was then applied unchanged to WT and disease cells in that stratum.

Module-high occupancy was summarized as the fraction of cells exceeding the matched WT threshold. The corresponding disease-associated change (ΔF) was defined as the disease-minus-WT difference in module-high fraction, with positive values indicating increased occupancy and negative values indicating reduced occupancy. Pooled-cell ΔF was used as the primary effect-size measure for visualization and comparison of cellular-state remodeling in Figures 2–3. Module-high fractions were additionally calculated independently for each biological sample for replicate-level analyses.

Continuous AUCell scores were retained for analyses of distributional displacement, width, and threshold dependence. Threshold sensitivity was evaluated using WT-derived q85, q90, and q95 thresholds within the same population-, age-, and module-matched strata.

#### 4.4 Statistical analysis of transcriptional-state changes

Transcriptional-state remodeling was evaluated using complementary cellular-distribution and biological-replicate analyses. Within each matched neuronal population, age, and module, changes in module-high occupancy were summarized by pooled-cell ΔF. Separation of WT and disease cellular-state distributions was evaluated using two-sided Fisher’s exact tests on module-high versus non-high cell counts, with Benjamini-Hochberg correction across modules within each neuronal population-age-genotype comparison. Adjusted values from these analyses are reported as cell-level FDR.

To evaluate reproducibility across biological samples, module-high fractions were also calculated independently for each sample after excluding sample–population combinations containing fewer than 10 cells. Genotype groups were compared using two-sided Wilcoxon rank-sum tests where biological replication permitted. Independent biological-replicate pseudobulk analysis of module-gene expression provided an additional assessment of transcriptional concordance, as described in Section 5.

#### 4.5 Detection-depth and transcript-complexity sensitivity analyses

To assess whether the principal AUCell effects were sensitive to transcript detection or library complexity, four complementary analyses were performed for Ex1_IT and Ex3_DeepCTCF neurons.

First, detected gene number (nFeature_RNA) and UMI count (nCount_RNA) were compared between Tau4RΔK-AP and age-matched WT cells after primary quality-control filtering.

Second, continuous E1 AUCell scores were analyzed with a linear mixed-effects model containing genotype, log-transformed UMI count, and detected gene number as fixed effects and biological-sample identity as a random intercept. Tau4RΔK-AP was the reference genotype.

Third, Tau4RΔK-AP and WT cells were matched 1:1 without replacement within each neuronal population and age by nearest-neighbor matching on detected gene number and UMI count, using a caliper of 0.20. Module-high occupancy was recalculated in the matched cells using the original WT-derived thresholds.

Fourth, raw count matrices were downsampled to 1,803 UMIs per cell, corresponding to the 10th percentile of UMI depth across the target populations. Gene rankings and AUCell scores were then recomputed using the same module definitions and scoring procedure as in the primary analysis.

These analyses were performed independently of the percentile-threshold sensitivity analyses.

### 5. Differential expression and pathway-level validation

Cell-level differential expression was used for neuronal marker identification, cluster annotation, and exploratory gene prioritization. Marker genes were identified in Seurat using FindAllMarkers with a Wilcoxon rank-sum test, a log-fold-change threshold of 0.5, and a minimum detected-cell fraction of 0.20.

For biological-replicate differential expression, raw counts were summed across cells from the same biological sample, neuronal population, and age to generate pseudobulk profiles. Pseudobulk analyses were performed with edgeR. Lowly expressed genes were filtered using filterByExpr, library-size normalization factors were estimated by trimmed mean of M-values normalization with calcNormFactors, and Tau4RΔK-AP and age-matched WT samples were compared separately within each neuronal population and age using quasi-likelihood generalized linear models fitted with glmQLFit and tested with glmQLFTest. Genotype was the model term and WT the reference group. Pseudobulk contrasts were evaluated only when both genotype groups contained at least two biological samples and each contributing sample contained at least 20 cells from the corresponding neuronal population.

To evaluate whether AUCell-defined occupancy changes were accompanied by coordinated differential expression of the corresponding module genes, each gene in a pseudobulk contrast was assigned the signed statistic sign(log₂FC_g) × min[−log₁₀(P_g), 50], where log₂FC_g and P_g are the pseudobulk log₂ fold change and nominal P value for gene g.

Module-level enrichment was summarized as the difference between the mean signed statistic among genes belonging to the module and the mean among all remaining genes in the same contrast. Enrichment was tested with two-sided Wilcoxon rank-sum tests comparing module and background genes, with Benjamini-Hochberg correction across modules within each neuronal population and age.

Concordance between cell-state remodeling and biological-replicate transcriptional change was assessed by comparing ΔF with the corresponding pseudobulk module-enrichment score across matched neuronal population, age, and module comparisons.

### 6. Statistical and Distributional Remodeling Analyses

#### 6.1 Distributional remodeling metrics

Continuous AUCell distributions were summarized within matched neuronal populations using four complementary quantities, all calculated on pooled cells. State-occupancy change was represented by ΔF as defined in Section 4.3. Central displacement was the disease-minus-WT difference in median AUCell score. Wasserstein-1 distance was calculated between the empirical disease and WT AUCell distributions and assigned the sign of the corresponding median displacement. Distributional width was summarized as the log2 ratio of disease to WT AUCell-score variance, with 10 added to the variances for numerical stability.

These quantities were interpreted jointly to distinguish changes in high-state occupancy, central displacement, broadening, and narrowing.

#### 6.2 Statistical testing of distributional differences

Cell-level tests were used to characterize complementary features of the AUCell distributions within matched neuronal populations. Module-high versus non-high frequencies were compared using two-sided Fisher’s exact tests; 95% Wilson score intervals were calculated where shown. Continuous AUCell scores were compared using two-sided Wilcoxon rank-sum tests, overall distributional differences using two-sample Kolmogorov-Smirnov tests, and dispersion using Fligner-Killeen tests.

Benjamini-Hochberg correction was applied separately within the corresponding Fisher, Wilcoxon, Kolmogorov-Smirnov, and Fligner-Killeen analysis families. Adjusted values from these pooled-cell distributional tests are cell-level FDR measures and were interpreted together with ΔF, median displacement, signed Wasserstein distance, and variance change.

#### 6.3 Threshold sensitivity and stability analyses

Sensitivity to the module-high definition was evaluated using alternative WT-derived percentile thresholds. Thresholds were calculated independently within the matched WT neuronal population, age, and module and then applied unchanged to the corresponding WT and disease cells.

Figure 2 used q85, q90, and q95 thresholds, whereas the Figure 3 distributional analysis used q80, q90, and q95. ΔF was recalculated at each threshold, and directional stability was defined as retention of the same sign across the evaluated thresholds.

#### 6.4 Integrated composition-state decoupling and remodeling analyses

Population representation and within-population transcriptional-state remodeling were evaluated as separate dimensions. Representation was summarized by population-level log2 fold change relative to age-matched WT, whereas module-specific state remodeling was summarized by ΔF.

For descriptive visualization in Figure 3A, |log2FC| < 0.5 was used to denote the absence of a major representation shift, and |ΔF| ≥ 0.05 was used to denote an appreciable occupancy change. These thresholds were used only to organize population-module combinations visually and did not define statistical significance.

Overall remodeling magnitude for each neuronal population and age was calculated as the mean absolute ΔF across the selected module set. The maximum absolute ΔF and the number of modules meeting the corresponding cell-level FDR criterion were retained as complementary measures of remodeling magnitude and breadth. Where categorical descriptors were required for visualization, representation was classified using the ±0.5 log2FC thresholds described above and remodeling magnitude relative to the median remodeling score across the analyzed population-age combinations. Representation and state remodeling were not combined into a single vulnerability score.

### 7. Single-cell ATAC-seq analysis

#### 7.1 Cohort and neuronal dataset construction

Previously generated cortical single-cell ATAC-seq data from the Tau4RΔK-AP disease-progression cohort were reanalyzed to assess whether selected RNA-defined transcriptional-state programs were accompanied by corresponding changes in chromatin accessibility. Cross-modal analyses focused on the 6- and 12-month cohorts.

Previously annotated neuronal nuclei were extracted from the cortical scATAC-seq dataset and analyzed using Signac and Seurat. ATAC neuronal populations were matched to the major RNA-defined neuronal classes using broad neuronal identity and marker profiles. Because the two modalities did not resolve identical cellular subdivisions, RNA-ATAC correspondence was defined at the level of broad biological identity rather than exact cluster equivalence.

The principal ATAC populations used for cross-modal analyses are referred to as aEx1, aEx2, aEx3, aIn1, and aIn2 in the corresponding figures.

#### 7.2 Chromatin preprocessing and neuronal annotation

Gene annotations were obtained from EnsDb.Mmusculus.v79, converted to UCSC-style chromosome notation, and assigned to the mm10 genome assembly. Chromatin-accessibility data were processed using Signac.

Gene-level accessibility estimates were generated using GeneActivity, and the resulting assay was log-normalized. Neuronal identities were assigned by integrating existing ATAC annotations with GeneActivity-based marker profiles and canonical excitatory- and inhibitory-neuronal identity markers.

A manually curated RNA-ATAC crosswalk was then generated on the basis of broad neuronal identity. These assignments were used solely for cross-modal comparison and were not interpreted as evidence that individual RNA and ATAC clusters represented identical transcriptomic subtypes.

#### 7.3 Pathway-associated GeneActivity scoring

Selected S-, E-, and R-family modules from the RNA analysis were transferred to the scATAC-seq GeneActivity matrix. Modules were analyzed individually rather than collapsed into family-level scores. Gene symbols from each module were matched to features represented in the GeneActivity assay, and modules with insufficient gene representation were excluded.

For each nucleus, module-associated GeneActivity was calculated as the mean normalized GeneActivity across represented genes belonging to that module. Scores were then summarized by neuronal population, age, genotype, biological sample, and module.

#### 7.4 Statistical analysis of pathway-associated accessibility

Disease-associated GeneActivity differences were evaluated separately within matched neuronal populations and ages. For cell-level analyses, module-associated GeneActivity distributions were compared between Tau4RΔK-AP and WT nuclei using two-sided Wilcoxon rank-sum tests where sufficient nuclei were available, and effect magnitude was summarized using the standardized mean difference. Benjamini-Hochberg correction was applied within the corresponding module-population-age analysis families.

Where biological replication permitted, GeneActivity scores were also summarized independently within biological samples before genotype comparison. Because later scATAC-seq groups contained limited biological replication, the principal cross-modal interpretation focused on the direction and magnitude of accessibility changes rather than animal-level genotype inference.

#### 7.5 RNA-ATAC cross-modal comparison

RNA and ATAC neuronal populations were linked using the broad-identity crosswalk described above. Cross-modal comparisons were restricted to population-module combinations with biologically interpretable counterparts in both modalities.

RNA remodeling was represented by ΔF, whereas ATAC remodeling was represented by the corresponding change in module-associated GeneActivity. Effects were considered directionally concordant when both modalities changed in the same direction and discordant when their directions differed. Because RNA and ATAC measurements were obtained from separate cells and GeneActivity does not directly measure an RNA-defined module-high state, accessibility changes were interpreted as cross-modal correlates of transcriptional remodeling rather than as direct evidence of a regulatory mechanism.

### 8. Independent TauP301S-AP single-nucleus RNA-sequencing cohort

#### 8.1 Nuclei isolation, library preparation, and sequencing

Single-nucleus RNA sequencing was performed independently on cerebral cortex and hypothalamus from 6-month-old male WT and TauP301S-AP mice. Nuclei were isolated from frozen brain tissue using an ice-cold nuclei-isolation workflow comprising mechanical homogenization, filtration, centrifugation, washing, and microscopic assessment before library preparation.

Single-nucleus libraries were generated using the Parse Biosciences Evercode WT Mini kit, version 3, according to the manufacturer’s protocol. Libraries were sequenced on an Illumina NovaSeq 6000 using an SP Reagent Kit v1.5, 100 cycles, targeting a sequencing depth of at least 30,000 reads per nucleus. Cortex and hypothalamus were processed and analyzed as separate regional datasets.

#### 8.2 Preprocessing, clustering, and neuronal annotation

Cortical and hypothalamic datasets were processed independently. Quality-controlled nuclear expression matrices were normalized, dimensionally reduced, and clustered separately for each brain region.

Initial neuronal identities were assigned using MapMyCells^33^ reference mapping and previous publications^32,34^ and subsequently evaluated using neurotransmitter identity, cortical-layer and projection markers, neuropeptides, transcription factors, and regionally informative marker expression. Final figure labels describe the dominant biological identity supported by these combined annotations. Reference labels were not interpreted as evidence of exact one-to-one equivalence with external atlas subtypes or with neuronal populations in the Tau4RΔK-AP discovery cohort.

Cortical populations included IT-like, corticothalamic, corticofugal, Rorb-positive, Car3-like, DG-like, and inhibitory neuronal identities. A D2 spiny projection neuron population was also recovered but was excluded from the principal cortical comparison framework. Hypothalamic populations encompassed arcuate, supramammillary, ventromedial hypothalamic, mammillary, zona-incerta-like, and tuberomammillary identities. The Hdc-positive population is referred to as TMN HDC and interpreted as a tuberomammillary histaminergic population.

#### 8.3 AUCell pathway-state scoring and statistical analysis

The fixed E-, S-, and R-family modules established in the Tau4RΔK-AP discovery analysis were applied unchanged to the cortical and hypothalamic TauP301S-AP datasets. Figure 5 used previously generated per-nucleus AUCell scores.

Module-high thresholds were defined separately for cortex and hypothalamus. For each module, the reference threshold was the 90th percentile of its AUCell distribution across all WT neuronal nuclei within the corresponding brain region. This regional WT threshold was applied unchanged to WT and TauP301S-AP nuclei from every neuronal population in that region. Because thresholds were defined across all WT neurons rather than separately within each population, WT module-high occupancy was not constrained to 10% within individual neuronal populations.

Within each neuronal population and module, WT and TauP301S-AP nuclei were compared directly using a two-sided Fisher’s exact test on module-high versus non-high counts. Comparisons required at least 20 nuclei from each genotype. Benjamini-Hochberg correction was applied across tested modules within each neuronal population. These adjusted values are referred to as cell-level FDR.

The effect size for each comparison was ΔF, calculated as the TauP301S-AP-minus-WT difference in module-high fraction. Positive values indicate increased and negative values reduced occupancy of the corresponding module-high state.

#### 8.4 Neuronal representation and transcriptional-state remodeling

Neuronal representation and transcriptional-state remodeling were analyzed as separate dimensions. Within cortex and hypothalamus, relative representation was calculated as the fraction of quality-controlled neuronal nuclei assigned to each population.

For visualization, TauP301S-AP-associated representation change was expressed as log2[(pTauP301S-AP + 10^-5^) / (pWT + 10^-5^)], where p denotes the observed genotype-level population fraction. The pseudocount was used only to stabilize visualization of low-frequency populations and did not represent an inferential adjustment.

Because the independent cohort was not designed for strongly powered animal-level composition inference, Figure 5 composition was interpreted descriptively from observed representation fractions and effect sizes. Pooled-nucleus counts were not used to establish genotype significance for relative representation.

Transcriptional-state remodeling was quantified from module-specific changes in module-high occupancy. For Figure 5D, conditional remodeling magnitude was defined as the mean absolute occupancy change among modules meeting the cell-level FDR criterion within the corresponding neuronal population. Populations with no module meeting this criterion were assigned a value of zero. For Figure 5E, overall remodeling magnitude was calculated as the mean absolute occupancy change across all tested E-, S-, and R-family modules, irrespective of statistical classification.

Remodeling breadth was defined as the number of tested modules meeting the cell-level FDR criterion. Because module eligibility depended on expression and minimum-cell requirements, the number of tested modules was retained separately for each neuronal population.

For the hypothalamic visualization, net direction was summarized from the signed occupancy effects of modules meeting the cell-level FDR criterion, weighted by their statistical support. Positive values indicate remodeling dominated by increased module-high occupancy in TauP301S-AP, whereas negative values indicate remodeling dominated by decreased occupancy.

### 9. Human Alzheimer’s disease analysis

#### 9.1 Human neuronal datasets

Human neuronal single-nucleus transcriptomic data were analyzed using pre-existing quality-controlled Seurat objects containing published high-resolution neuronal annotations and donor metadata^5^. Analyses were restricted to prefrontal cortex (brainRegion == “PFC”).

Four neuronal datasets were analyzed independently: three excitatory-neuron datasets (ExcSet1-ExcSet3) and one inhibitory-neuron dataset (Inhibitory). Metadata used for downstream analyses included neuropathological disease-stage classification (ADdiag3types), donor identifier (subject), sex (msex), and published high-resolution neuronal subtype (cell_type_high_resolution). Donors were classified as nonAD, earlyAD, or lateAD. Published neuronal-subtype labels were retained directly rather than reclustering human neurons according to mouse-derived boundaries.

#### 9.2 Cross-species pathway scoring

The fixed human gene sets corresponding to the E-, S-, and R-family modules established in the mouse analysis were applied to the human datasets. Module scoring was performed independently within each source dataset using AUCell and log-normalized RNA expression.

Per-cell gene rankings were generated with AUCell_buildRankings, and continuous module scores were calculated with AUCell_calcAUC. A module was retained when at least 10 genes from its finalized definition were represented in the corresponding expression matrix. AUCell scores were linked to donor, disease stage, sex, source dataset, and published neuronal-subtype metadata.

Cross-species comparisons were made between biologically related neuronal classes and did not assume one-to-one equivalence between mouse and human transcriptomic subtypes.

#### 9.3 Donor-balanced definition of human module-high states

Module-high states were defined using non-AD donors as the reference group. To prevent donors contributing large numbers of nuclei from dominating threshold estimation, the q90 AUCell threshold was calculated separately within each eligible non-AD donor for each neuronal subtype and module. Only donors contributing at least 20 nuclei to the corresponding subtype were included.

The final reference threshold was the median of these donor-specific q90 values and was applied unchanged to nuclei from non-AD, early-AD, and late-AD donors of the corresponding neuronal subtype. Module-high occupancy was then calculated independently for each donor, subtype, and module.

Disease-stage occupancy effects were summarized as differences in mean donor-level module-high fraction for early AD versus non-AD, late AD versus non-AD, and late AD versus early AD.

#### 9.4 Donor-level statistical inference

Donors were the unit of biological inference for the primary human analyses. Donors contributing fewer than 20 nuclei to a neuronal subtype were excluded, and subtype-module analyses required at least eight eligible donors in each disease-stage group. Continuous AUCell scores were summarized within each donor-neuronal subtype-module combination before disease-stage testing.

Disease-stage effects were evaluated with linear mixed-effects models of donor-mean AUCell score on disease stage, sex, age at death, postmortem interval, and scaled median detected gene number, with batch as a random intercept and nonAD as the reference stage. LateAD-versus-earlyAD contrasts were obtained by releveling the disease-stage factor. Covariate adjustment was restricted to variables available in the released donor metadata.

Donor-level changes in module-high occupancy were retained as effect-size measures describing redistribution into or out of the high-state tail.

#### 9.5 Cell-level sensitivity analyses

Pooled-cell analyses were used only as secondary sensitivity analyses of the underlying cellular distributions. Within each neuronal subtype and disease-stage contrast, module-high versus non-high frequencies were compared using two-sided Fisher’s exact tests, and continuous per-cell AUCell scores were compared using Wilcoxon rank-sum tests where indicated. Benjamini-Hochberg correction was applied within the corresponding analysis families.

#### 9.6 Ranked human neuronal remodeling burden

Aggregate remodeling was compared across human neuronal subtypes using the selected E-, S-, and R-family modules represented in the principal cross-species analysis.

The primary score determining the rank order in Figure 4E was the unweighted remodeling score, defined as the mean absolute donor-level late-AD-versus-non-AD difference in module-high occupancy across evaluated modules. Rank 1 therefore denotes the neuronal population with the largest mean absolute remodeling across the selected module panel.

Weighted and signed versions of the remodeling score were calculated as complementary sensitivity measures. The weighted score incorporated donor-level statistical support, whereas the signed score retained effect direction and distinguished populations dominated by increases from those dominated by decreases. Neither measure determined the primary rank order shown in Figure 4E. These scores summarize transcriptional-state remodeling and were not interpreted as measures of neuronal depletion.

#### 9.7 Leave-one-donor-out ranking sensitivity

Sensitivity of the Figure 4E ranking to individual donors was assessed by repeating the ranking analysis after excluding one donor at a time. For each iteration, donor-level disease-stage effects and the unweighted remodeling score were recalculated using the remaining eligible donors, and neuronal populations were reranked.

The range of ranks obtained across leave-one-donor-out iterations is shown in Figure 4E. This analysis was used to determine whether the relative ordering of neuronal populations was driven disproportionately by individual donors.

### 10. Cross-dataset definition of neuronal state architecture

Neuronal involvement was evaluated using three analytical dimensions quantified separately within each dataset: population representation, transcriptional-program remodeling within matched neuronal identities, and remodeling of continuous cellular-state distributions. These dimensions were analyzed separately and were not combined into a composite vulnerability score.

Cross-dataset comparisons were performed at the level of broad neuronal identity and pathway-associated state rather than requiring one-to-one correspondence between transcriptomic clusters. Correspondence among the Tau4RΔK-AP discovery cohort, the independent TauP301S-AP cohort, and human neuronal populations was assigned on the basis of neuronal class, regional identity, and established marker expression. These mappings were interpreted as biological correspondences rather than exact transcriptomic equivalence.

Cross-dataset recurrence was evaluated at two levels. Architectural recurrence referred to reappearance of higher-order features, including remodeling within related neuronal classes, engagement of the same E-, S-, or R-family biological programs, and dissociation between relative representation and within-population molecular state. Architectural recurrence did not require identical pathway effects or effect magnitudes.

Directional recurrence was assessed separately for individual pathways and required effects to change in the same direction across the compared datasets. Effects in opposite directions were treated as directional divergence even when they involved the same pathway or broader biological program.

The same framework was used for mouse-human comparisons. Human neuronal populations were compared with biologically related mouse identities and pathway-state programs without assuming one-to-one subtype homology. Chromatin-accessibility analyses were treated as cross-modal supporting evidence rather than as a required component of the transcriptional-state framework.

### 11. Neuro2a cell culture, pharmacological perturbation, and bulk RNA sequencing

#### 11.1 Neuro2a cell culture and retinoic acid conditioning

Mouse Neuro2a neuroblastoma cells were obtained from the research group of Olav Michael Andersen. Cells were maintained in T75 culture flasks in high-glucose Dulbecco’s modified Eagle medium (DMEM; Gibco, cat. no. 41966-029) supplemented with 10% fetal bovine serum and 1% penicillin-streptomycin at 37°C in a humidified atmosphere containing 5% CO .

For experiments, cells were seeded at (2.0 x 10^4^) cells per well in 24-well plates. Plates were coated with poly-D-lysine before seeding. Following incubation for 60 min at room temperature, coating solution was removed, wells were washed three times with cell-culture-grade water, and plates were allowed to dry under sterile conditions before use.

Cells were detached from maintenance cultures using TrypLE, collected in complete growth medium, centrifuged at 300 × (g) for 5 min, and resuspended in fresh growth medium. Cell concentration and viability were assessed by trypan-blue exclusion before seeding.

Cells were seeded on day 0 in high-glucose DMEM containing 10% fetal bovine serum and 1% penicillin-streptomycin. On day 1, medium was replaced with RA-conditioning medium consisting of high-glucose DMEM supplemented with 2% fetal bovine serum, 1% penicillin-streptomycin, and 30 µM all-trans retinoic acid (Sigma-Aldrich, cat. no. R2625). Retinoic acid was prepared from a 100 mM stock in dimethyl sulfoxide (DMSO), resulting in a final DMSO concentration of 0.03%. RA-containing solutions were protected from light during preparation and handling. Cells were maintained under RA-conditioning conditions until pharmacological treatment on day 6, with medium replaced every other day.

#### 11.2 Single-molecule fluorescent *in situ* hybridization (smFISH)

smfISH was performed in cultured cells as previously described^35^, with minor modifications to accommodate the transcript targets analyzed in the present study. Briefly, gene-specific probe sets were designed against the relevant human or mouse transcripts using OligoMiner-derived resources and custom filtering, and 24-50 exon-targeting probes were selected per transcript. Probes were extended by primer-exchange reaction and hybridized to fixed and permeabilized cells, followed by hybridization of fluorescent oligonucleotides complementary to the corresponding barcodes. Images were acquired on an Olympus APEXVIEW APX100 fluorescence microscope using a 40× objective, tiled XY acquisition, and z-stacks, with identical regions re-imaged across successive hybridization rounds. Probe identities, target genes, PER barcodes, and fluorophores used in this study are provided in Table S19.

#### 11.3 Pharmacological pretreatment and oxidative-stress challenge

On day 6, RA-conditioned Neuro2a cells were pretreated for 2 h with 400 nM MK-8722, 40 µM AICAR, 400 nM PD0325901, or the corresponding vehicle. Drug stocks and working solutions were prepared according to the solvent conditions specified for each compound, and vehicle exposure was matched between the relevant treatment and control conditions.

Following the 2-h pretreatment period, cells received either hydrogen peroxide to a final concentration of 500 µM or the corresponding vehicle without replacement of the pretreatment medium. Cells were then incubated for a further 24 h.

The viability-associated experiment included the following conditions: vehicle, H O, MK-8722, AICAR, PD0325901, H O + MK-8722, H O + AICAR, and H O + PD0325901, together with cell-free background wells.

The bulk RNA-sequencing experiment used a separate four-condition factorial design comprising vehicle, H O, MK-8722, and H O + MK-8722. AICAR and PD0325901 were not included in the bulk RNA-sequencing analysis.

#### 11.4 PrestoBlue metabolic-activity assay

Cellular reducing activity was measured after 24 h of H O exposure using PrestoBlue HS Cell Viability Reagent (Thermo Fisher Scientific/Invitrogen, cat. no. P50200).

Treatment medium was replaced with prewarmed assay medium containing PrestoBlue HS diluted 1:10 in FluoroBrite medium supplemented with 10% fetal bovine serum and 1% penicillin-streptomycin. Cells were incubated at 37°C before fluorescence measurement using a VICTOR Nivo multimode plate reader with 540/30-nm excitation and 580/20-nm emission filters. Cell-free wells were included to estimate assay background.

Fluorescence values were background-corrected using the corresponding cell-free wells and expressed relative to the mean vehicle-control signal from the same experiment. Because PrestoBlue measures cellular reducing activity and can be influenced by metabolic state as well as viable cell number, it was interpreted as a viability-associated metabolic readout. Condition-level effects were summarized descriptively.

#### 11.5 Collection of samples for bulk RNA sequencing

For the four-condition bulk RNA-sequencing experiment, cells were collected after the 24-h oxidative-stress period. Treatment medium was removed, cells were washed once with room-temperature phosphate-buffered saline, and cells were detached using 300 µL TrypLE per well for 5 min at 37°C. TrypLE was neutralized with 500 µL high-glucose DMEM containing 10% fetal bovine serum and 1% penicillin-streptomycin.

Each biological replicate was collected separately and centrifuged at 300 × (g) for 5 min. Supernatant was removed, and the resulting cell pellet was resuspended in 60 µL 1× DNA/RNA Shield. Lysates were mixed thoroughly and stored frozen until RNA sequencing.

Bulk RNA sequencing was performed by Plasmidsaurus using Illumina sequencing, targeting approximately 10 million deduplicated reads per sample. Gene-level raw count matrices and sequencing quality-control outputs were used for downstream analyses.

#### 11.6 Bulk RNA-sequencing processing and differential-expression analysis

Bulk RNA-sequencing analysis included vehicle, H O, MK-8722, and H O + MK-8722 conditions. Ensembl identifiers were mapped to mouse gene symbols and annotations using org.Mm.eg.db, and lowly expressed genes were removed using the prespecified count filter in the analysis pipeline.

Differential expression was analyzed with DESeq2 using a full factorial design containing the main effects of H O and MK-8722 and their interaction. Six prespecified contrasts were evaluated: H O versus vehicle; MK-8722 versus vehicle; H O + MK-8722 versus H O ; H O + MK-8722 versus MK-8722; the H O × MK-8722 interaction; and H O + MK-8722 versus vehicle.

Hypothesis testing used DESeq2 Wald statistics with Benjamini-Hochberg FDR correction. Raw counts were used for differential-expression modeling. Variance-stabilized expression values were used only for sample-level visualization, including principal-component and sample-distance analyses.

#### 11.7 Bulk RNA pathway and E1-associated transcriptional scoring

The fixed module definitions established in the neuronal analyses were transferred to the Neuro2a bulk RNA-sequencing dataset to assess whether the corresponding transcriptional programs were pharmacologically modifiable. Because bulk RNA sequencing does not provide per-cell rankings, the single-cell AUCell/module-high framework was not applied.

Expression was transformed as log (CPM + 1) and standardized gene-wise across samples (z-score using the mean and population standard deviation across analyzed samples). The sample-level module score was the mean standardized expression across represented module genes. Because bulk RNA sequencing does not provide per-cell rankings, the AUCell/module-high framework was not applied, and the bulk score is a distinct quantity from ΔF despite using the same fixed gene sets.

The primary targeted analysis evaluated E1, derived from the NMDAR-linked AMPK signaling program. The bulk-RNA E1 score was interpreted as a transcriptional summary of the fixed E1 gene set and not as a direct measure of NMDAR signaling, CAMKK2 activity, or AMPK enzymatic activation.

## Data availability

Sequencing data generated in this study are being deposited in the ArrayExpress collection at EMBL-EBI BioStudies. Some sequencing data are available in the Gene Expression Omnibus under accession GSE175546. The datasets will be released publicly before publication.

## Code availability

Custom computer code and analysis scripts used in this study are available at https://github.com/thomaskim-lab/Vulnerability.

## Author contributions

S.S. and D.W.K. conceived the study. All authors contributed to the experimental work. S.S., M.M.P., and D.W.K. performed data analysis. D.W.K. supervised the study, acquired funding, and administered the project. S.S and D.W.K. wrote the manuscript.

## Acknowledgements

Some of the computing for this project was performed on the GenomeDK cluster. We thank GenomeDK and Aarhus University for providing computational resources and support that contributed to these research results. The authors gratefully acknowledge the technical and computational expertise and support provided by the Neuro Single Cell Platform (NeuSiC; Biotech Research & Innovation Centre [BRIC], University of Copenhagen). We specifically thank Irina Korshunova, Laura Wolbeck, and Eman Ahmad Mouhammad at NeuSiC for their valuable contributions.

We also thank Olav Michael Andersen’s group and Elnaz Fazeli for providing the Neuro2A cells. We further thank Thomas Willnow’s group and Karen-Marie Pedersen for their assistance and for providing access to the Victor Nivo plate reader.

## Supplemental Figure legends

**Figure S1.**
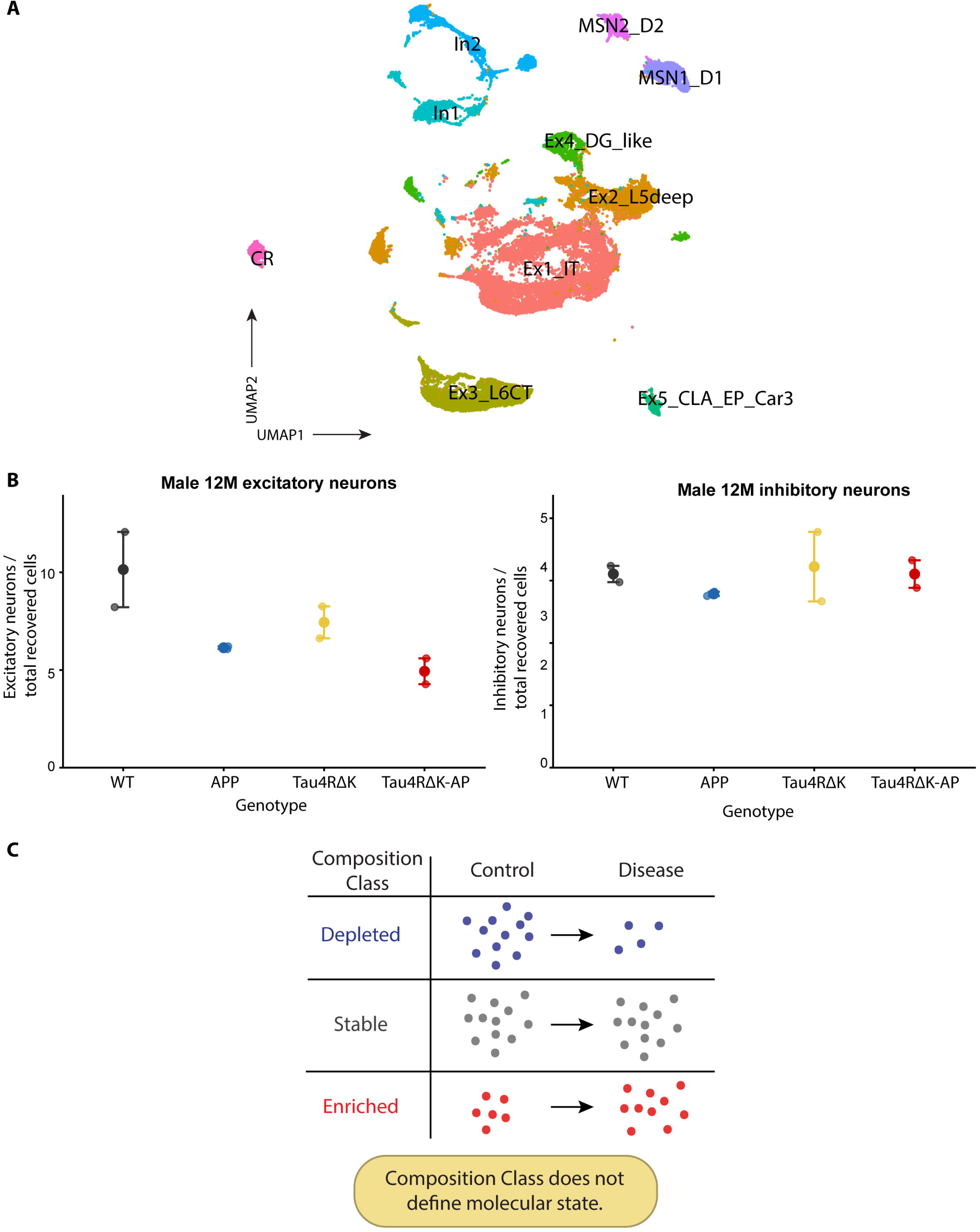
Neuronal population annotation, complementary male-cohort representation, and conceptual composition classes. **(A)** UMAP of neuronal cells from the Tau4RΔK-AP discovery dataset showing annotated neuronal identities. Principal populations used for composition analysis include Ex1_IT, Ex2_L5deep, Ex3_L6CT, Ex4_DG_like, In1_CGE_Calb2, and In2_MGE_CGE_mix. Additional annotated populations include Ex5_CLA_EP_Car3, D1 and D2 striatal spiny projection neurons, and Cajal-Retzius cells. **(B)** Relative representation of excitatory and inhibitory neurons in the complementary 12-month male cohort. Values are shown as the fraction of all quality-controlled cortical cells recovered from WT, APP;PS1, Tau4RΔK, and Tau4RΔK-AP mice. Individual points represent biological samples and error bars show mean ± SEM. **(C)** Conceptual classification of neuronal populations according to relative representation. Populations can show reduced, comparatively stable, or increased representation in disease relative to control. These categories describe relative recovery and do not define the molecular state of the neurons that remain or imply absolute neuronal loss.

**Figure S2.**
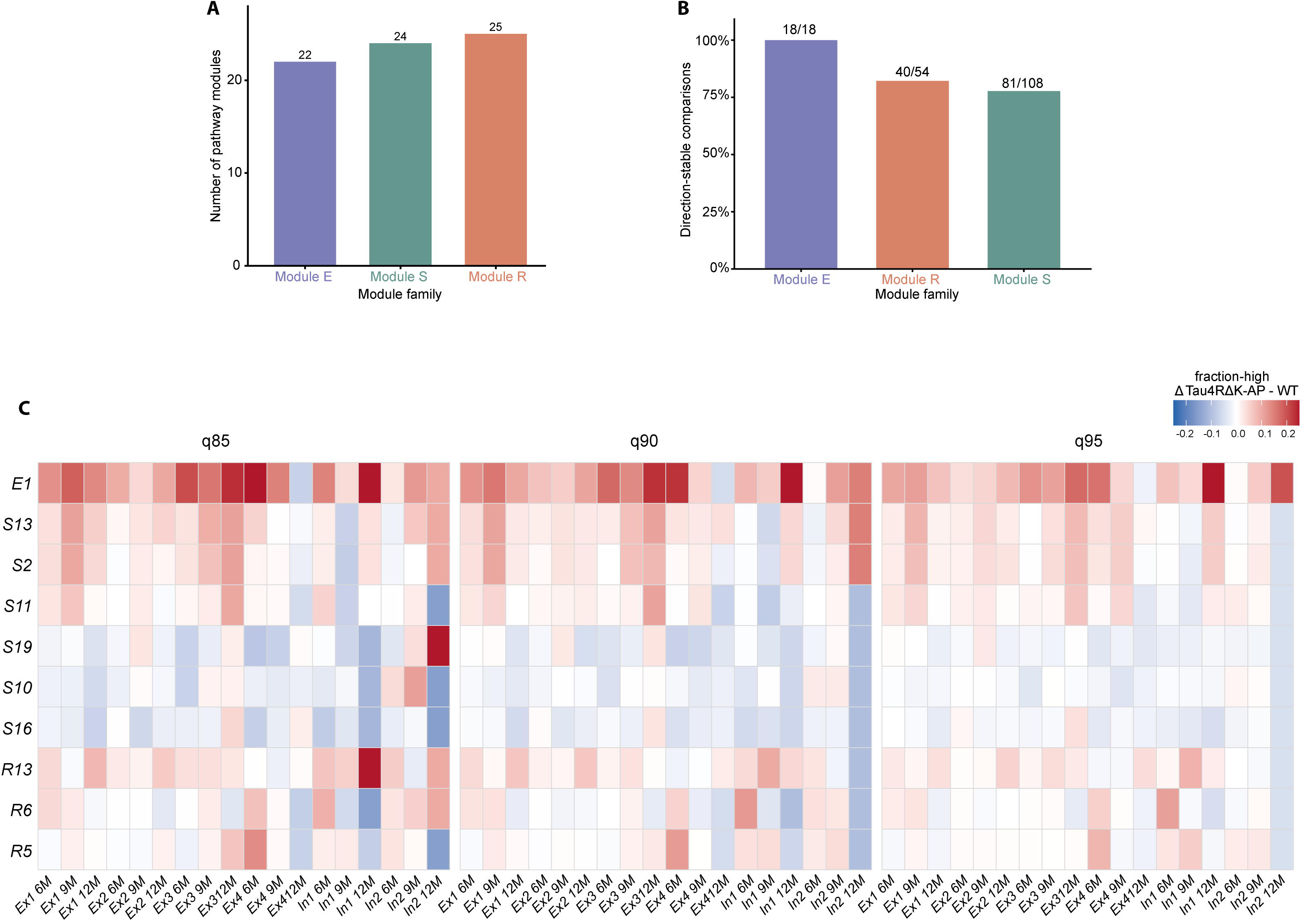
Module-family composition and robustness of state-remodeling direction to alternative AUCell thresholds. **(A)** Number of retained Reactome pathway modules assigned to each biological family after filtering and curation: 22 excitability, calcium, and ion-signaling modules (E), 24 synaptic-state modules (S), and 25 regulatory and stress-associated modules (R), for a total of 71 modules. **(B)** Directional stability of disease-associated module-state effects across alternative WT-derived AUCell thresholds. Bars show the proportion of comparisons retaining the same direction across q85, q90, and q95 thresholds. Values above bars indicate stable comparisons relative to the total evaluated: 18/18 for E-family comparisons, 40/54 for R-family comparisons, and 81/108 for S-family comparisons. **(C)** Threshold sensitivity of selected Tau4RΔK-AP-associated state effects across q85, q90, and q95 definitions of the module-high state. Rows show selected E-, S-, and R-family modules and columns show neuronal population by age combinations across Ex1_IT, Ex2_L5deep, Ex3_L6CT, Ex4_DG_like, In1_CGE_Calb2, and In2_MGE_CGE_mix. Color represents the Tau4RΔK-AP-minus-WT difference in module-high fraction, with red indicating increased and blue reduced occupancy.

**Figure S3.**
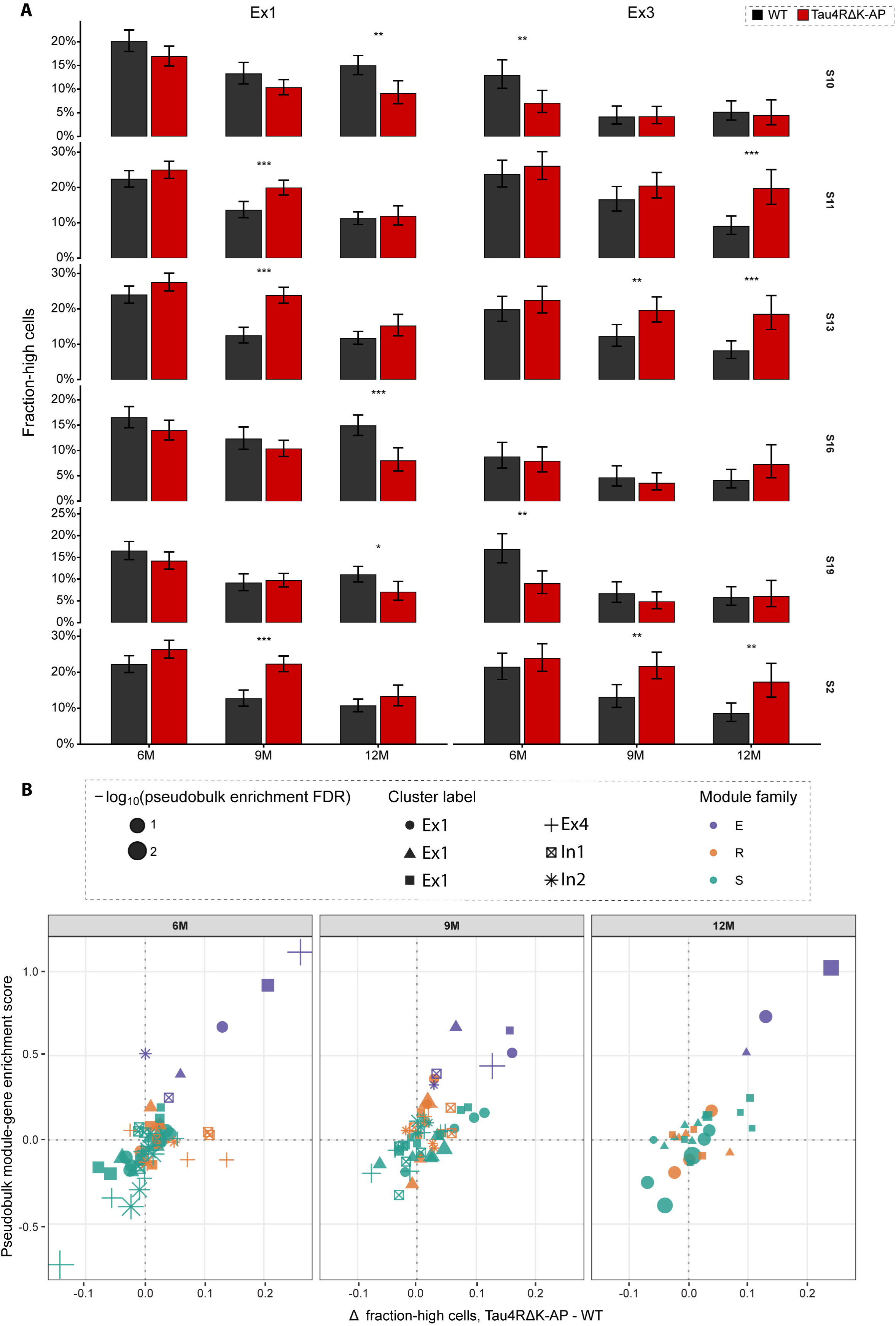
Synaptic-state remodeling and biological-replicate transcriptional support. **(A)** Module-high fractions for selected synaptic programs in Ex1_IT and Ex3_L6CT neurons from WT and Tau4RΔK-AP mice at 6, 9, and 12 months. Rows show S10, S11, S13, S16, S19, and S2. Bars show pooled cellular fractions and error bars show 95% Wilson score intervals. The selected programs show distinct and, in several comparisons, opposing trajectories within the same neuronal identities. **(B)** Relationship between AUCell-defined occupancy changes and sample-resolved pseudobulk module-gene enrichment at 6, 9, and 12 months. Each point represents a module within a matched neuronal population and age. The x-axis shows the Tau4RΔK-AP-minus-WT difference in module-high fraction and the y-axis shows the corresponding pseudobulk module-gene enrichment score. Point color denotes module family and point shape denotes neuronal identity. Point size represents the statistical support of the pseudobulk enrichment analysis. Dashed lines indicate zero change in each measurement.

**Figure S4.**
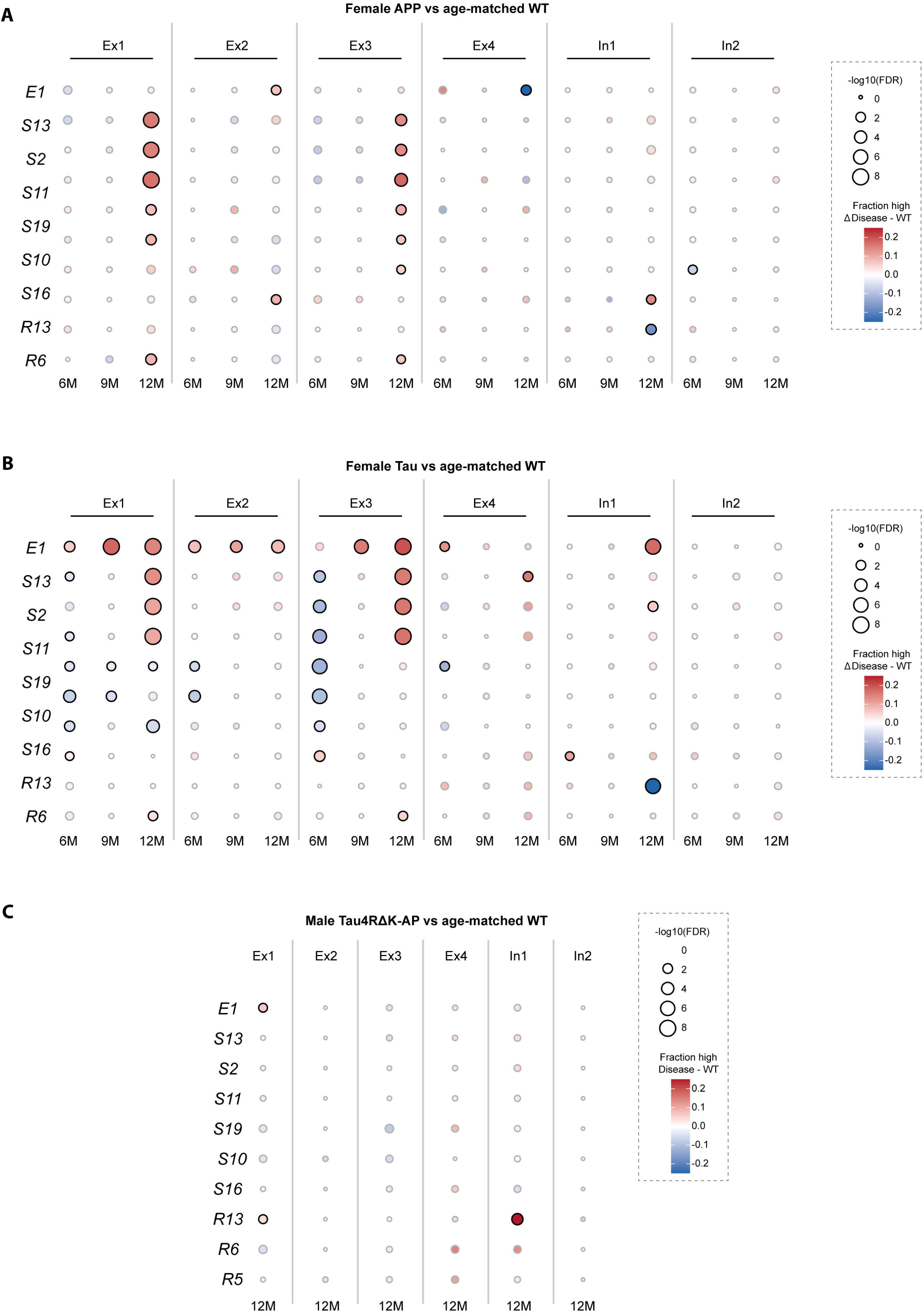
Genotype and sex context shape neuronal module-state remodeling. **(A)** Selected module-state effects in female APP;PS1 mice relative to age-matched WT at 6, 9, and 12 months across Ex1_IT, Ex2_L5deep, Ex3_L6CT, Ex4_DG_like, In1_CGE_Calb2, and In2_MGE_CGE_mix. **(B)** Corresponding module-state effects in female Tau4RΔK mice relative to age-matched WT. For **A** and **B**, rows show selected E-, S-, and R-family modules and columns show neuronal population by age combinations. Circle color represents the disease-minus-WT difference in module-high fraction, with red indicating increased and blue reduced occupancy. Circle size represents absolute effect magnitude, |Δfraction-high|. **(C)** Selected module-state effects in 12-month male Tau4RΔK-AP mice relative to age-matched male WT. Encoding is as in A and B.

**Figure S5.**
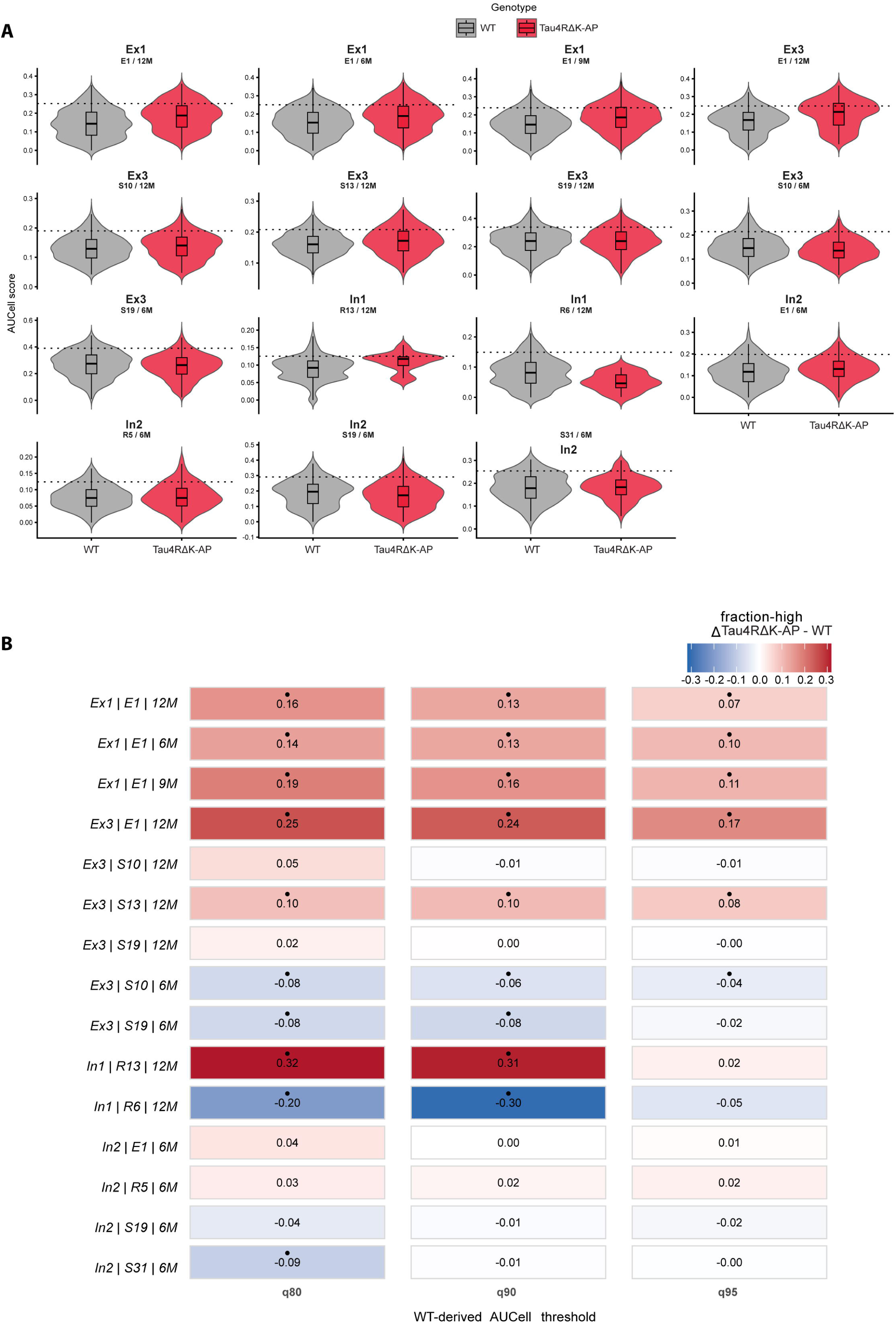
Distributional geometry and threshold sensitivity of selected neuronal state changes. **(A)** Per-cell AUCell-score distributions for selected neuronal population, module, and age combinations in WT and Tau4RΔK-AP mice. Violin plots show complete cellular score distributions, with embedded boxplots summarizing location and spread. The horizontal dotted line indicates the matched WT-derived q90 threshold used to define the module-high state. Examples include recurrent E1 remodeling in Ex1_IT and Ex3_L6CT, opposing synaptic-state changes in Ex3_L6CT, early state effects in In2_MGE_CGE_mix, and late R13 and R6 regulatory remodeling in In1_CGE_Calb2. **(B)** Sensitivity of selected disease-associated occupancy effects to q80, q90, and q95 WT-derived thresholds. Rows denote neuronal population, module, and age combinations. Cell color and printed values show the Tau4RΔK-AP-minus-WT difference in module-high fraction. Red indicates increased and blue reduced occupancy. Recurrent E1 effects retain their direction across thresholds, whereas selected regulatory and synaptic effects show stronger dependence on threshold placement.

**Figure S6.**
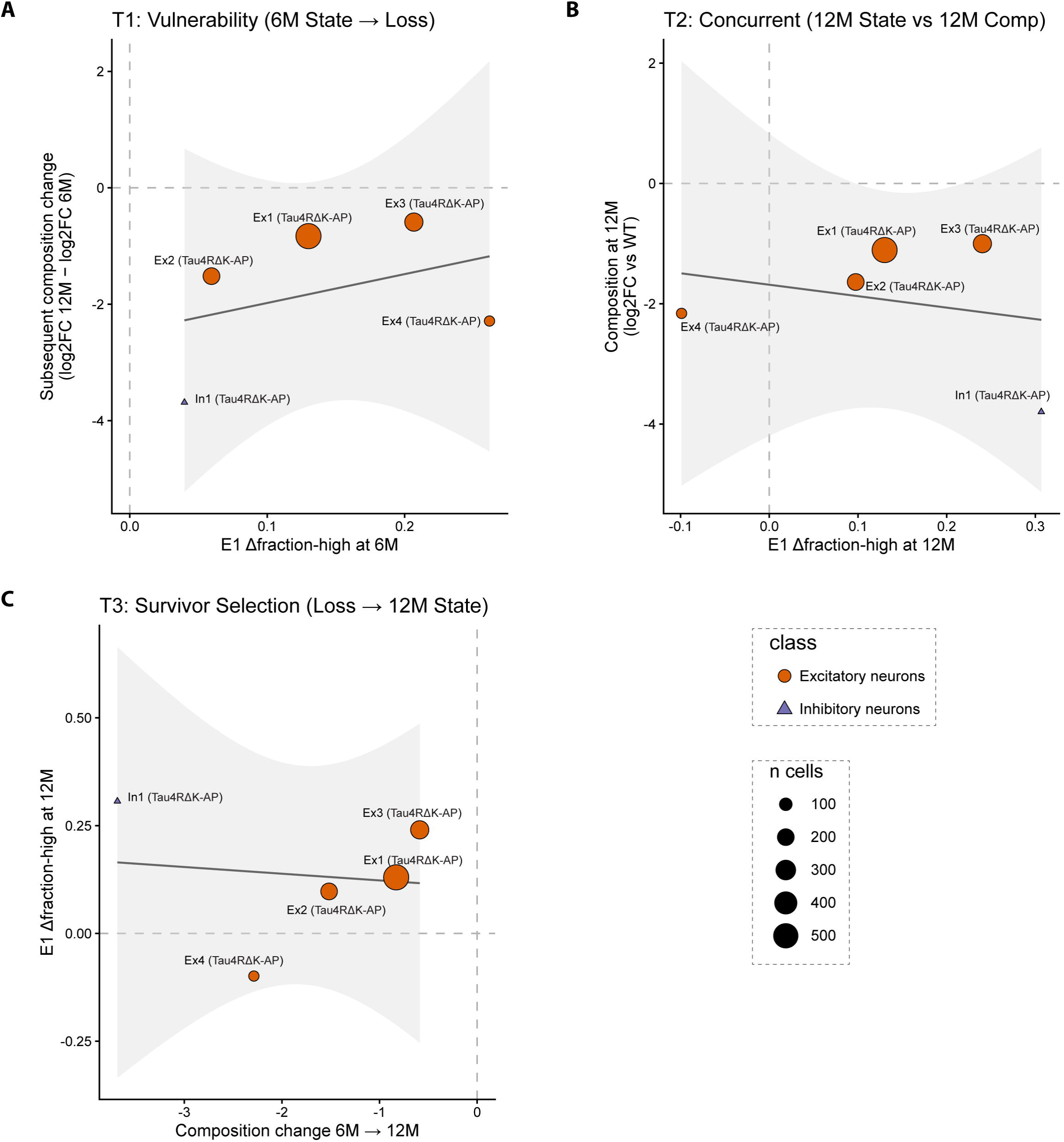
Early and late E1 states do not order age-associated changes in relative neuronal representation. **(A)** E1 occupancy change at 6 months plotted against the change in relative representation between the 6- and 12-month Tau4RΔK-AP cohorts for the neuronal populations included in the analysis. **(B)** E1 occupancy change at 12 months plotted against relative neuronal representation at 12 months. **(C)** Change in relative representation between 6 and 12 months plotted against E1 occupancy change among neurons recovered at 12 months. Points denote neuronal populations, with shape indicating excitatory or inhibitory identity and point size indicating the number of cells contributing to the corresponding state estimate. Lines show descriptive linear fits with 95% confidence intervals.

**Figure S7.**
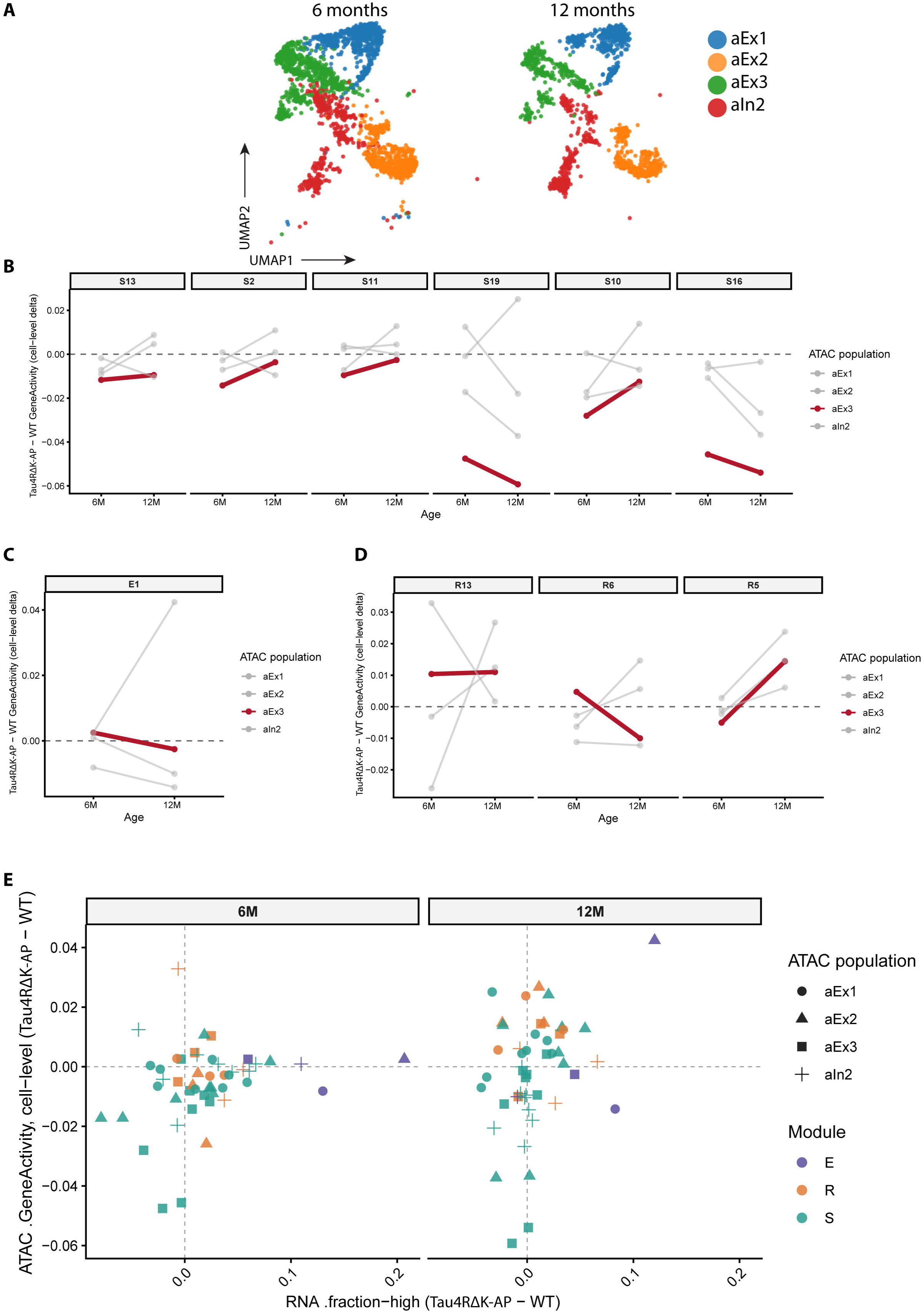
RNA-defined neuronal state remodeling shows heterogeneous chromatin-accessibility correlates. **(A)** UMAP representations of cortical scATAC-seq neuronal nuclei at 6 and 12 months. Nuclei are colored by ATAC-defined neuronal population. **(B)** Tau4RΔK-AP-minus-WT GeneActivity differences for selected synaptic-state modules S13, S2, S11, S19, S10, and S16 across ATAC neuronal populations at 6 and 12 months. Lines connect age-specific effects within populations. aEx3 is highlighted and other ATAC populations are shown for comparison. The dashed horizontal line indicates no GeneActivity difference. **(C)** Tau4RΔK-AP-minus-WT GeneActivity differences for E1 across ATAC neuronal populations at 6 and 12 months. **(D)** GeneActivity differences for regulatory programs R13, R6, and R5. Regulatory accessibility effects are similarly dependent on neuronal population and age. **(E)** Cross-modal comparison of RNA module-high occupancy changes and ATAC GeneActivity changes. Each point represents a matched population-module comparison. The x-axis shows the RNA Tau4RΔK-AP-minus-WT Δfraction-high and the y-axis shows the corresponding ATAC GeneActivity difference. Point shape denotes ATAC population and color denotes module family. Dashed lines indicate zero change.

**Figure S8.**
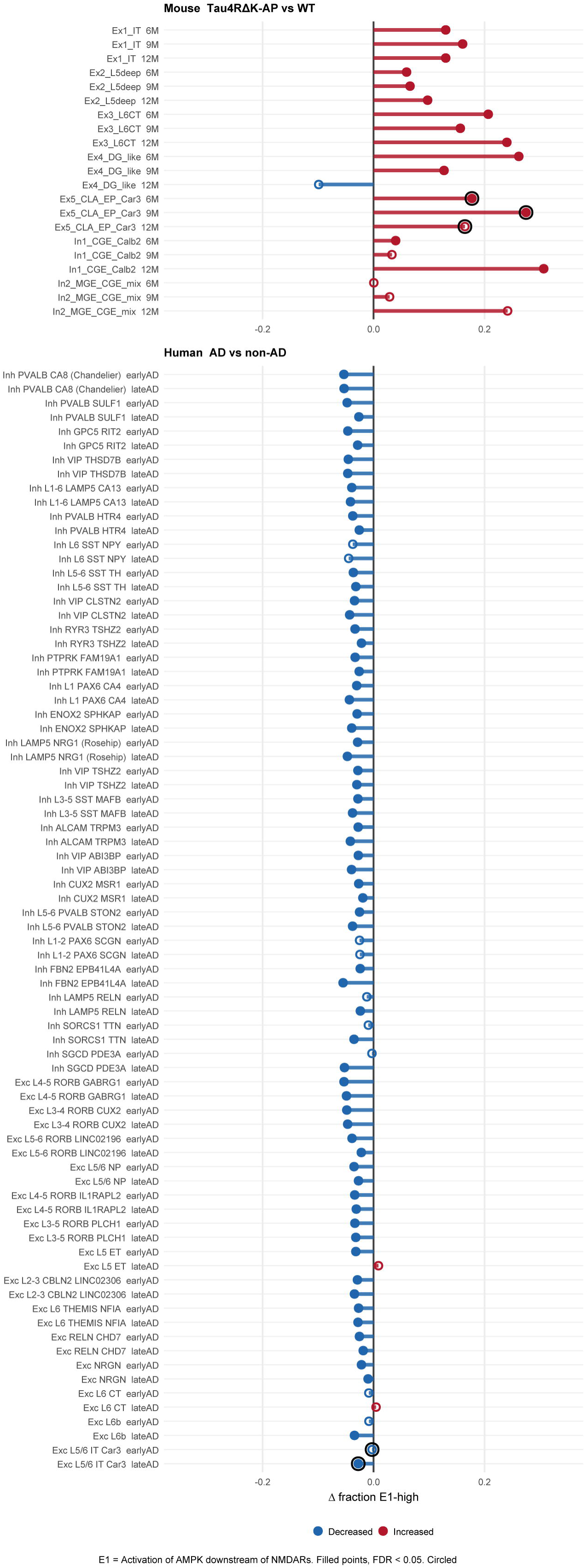
E1 remodeling differs in direction across mouse and human neuronal populations. The upper panel shows Tau4RΔK-AP-minus-WT differences in E1-high fraction across mouse neuronal populations and ages. Positive values indicate expansion and negative values depletion of the E1-high state. The mouse analysis demonstrates recurrent E1 expansion across glutamatergic identities at 6 and 9 months, together with population- and age-dependent divergence at later stages. The lower panel shows AD-minus-nonAD differences in E1-high occupancy across the broader set of human neuronal populations. EarlyAD and lateAD effects are displayed separately. Negative values predominate across human excitatory and inhibitory identities, in contrast to the recurrent glutamatergic E1 expansion in Tau4RΔK-AP mice. Filled mouse points denote comparisons meeting the cell-level FDR criterion used to characterize cellular-state separation in the discovery dataset. Filled human points denote comparisons meeting the donor-level FDR criterion, with donor as the unit of biological inference.

**Figure S9.**
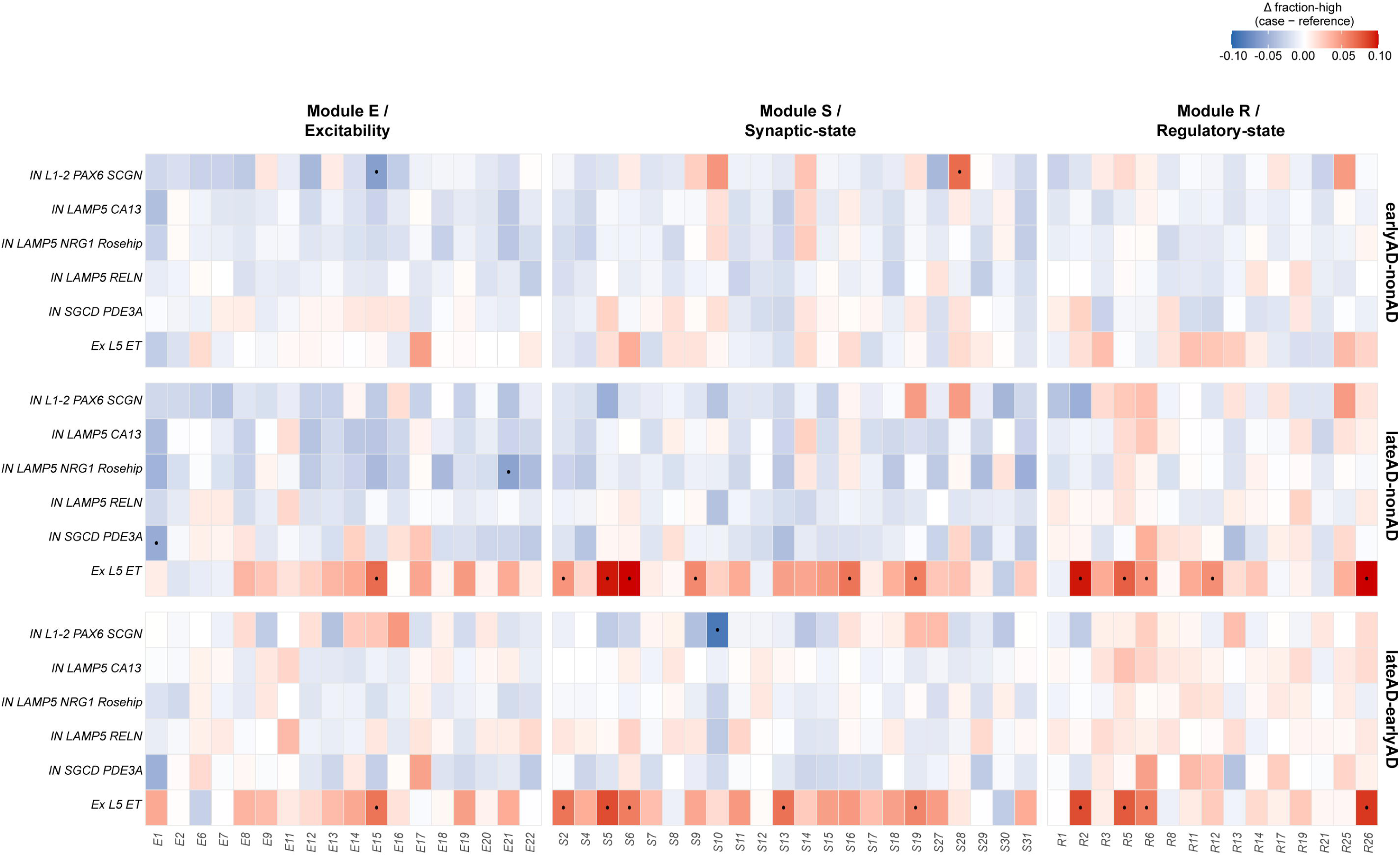
Mouse-derived state programs show selective engagement across human comparator populations. Heatmaps show disease-stage-associated differences in module-high occupancy across selected human neuronal populations used as comparator identities, together with EX L5 ET as a disease-responsive excitatory reference. Rows include L1/CGE-, LAMP5-, Rosehip-, and SGCD/PDE3A-related populations. Columns are grouped into excitability and calcium-signaling (E), synaptic-state (S), and regulatory and stress-associated (R) module families. Separate blocks show earlyAD versus nonAD, lateAD versus nonAD, and lateAD versus earlyAD. Cell color represents the case-minus-reference difference in donor-level module-high occupancy, with red indicating increased and blue reduced occupancy. Black dots denote donor-level FDR < 0.05. LAMP5 CA13 and LAMP5 RELN show comparatively limited remodeling, whereas the human-specific LAMP5 NRG1 Rosehip population shows a distinct pattern dominated by decreases. EX L5 ET shows broader lateAD-associated remodeling, illustrating the strong dependence of program engagement on neuronal identity.

**Figure S10.**
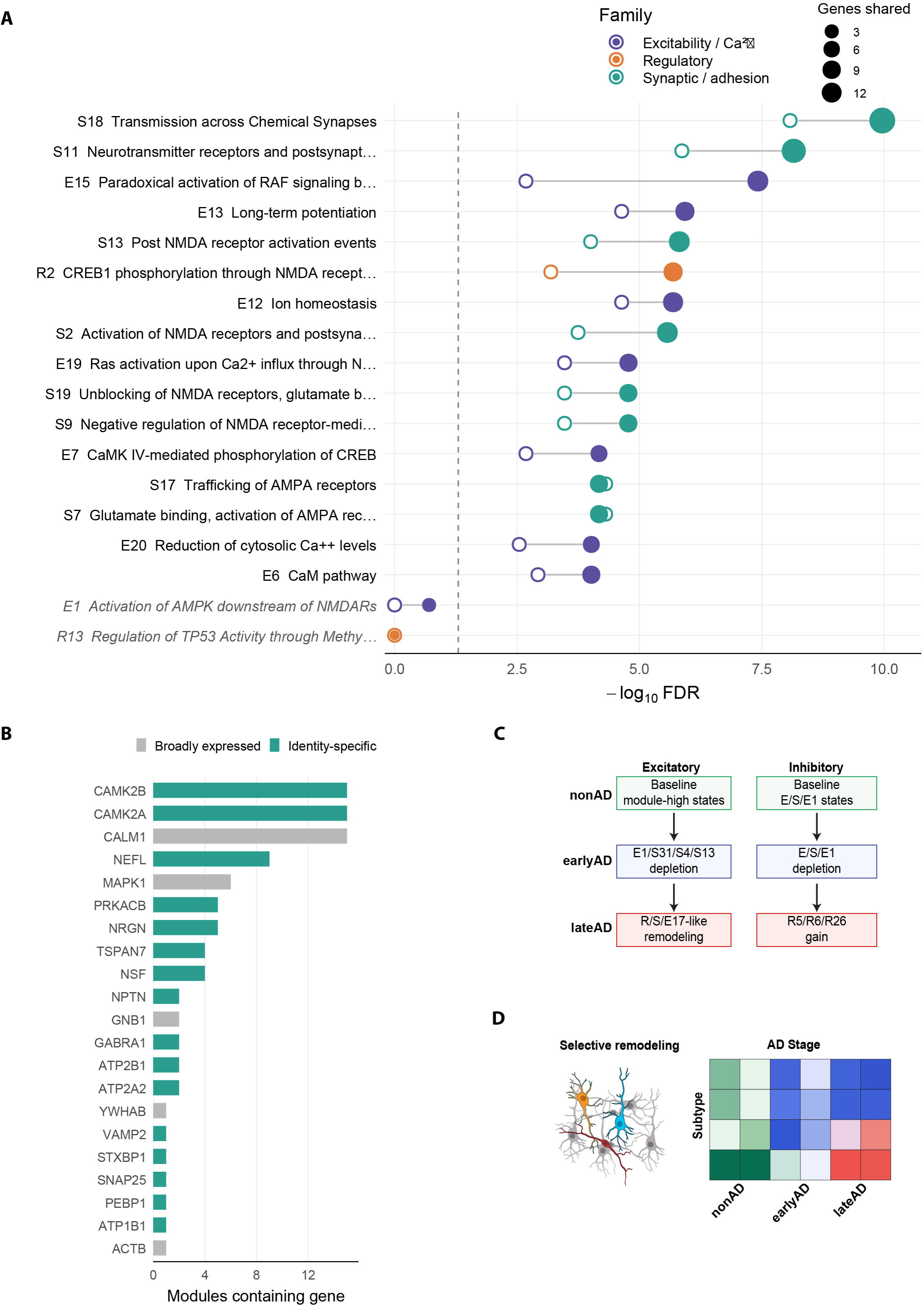
A control-derived human depletion-associated identity program overlaps selected synaptic and excitability-associated modules. **(A)** Enrichment of the independently defined human excitatory identity program^8^ across the fixed mouse-derived module set. The x-axis shows -log10 FDR and point size indicates the number of shared genes. Point color denotes module family. Paired symbols show enrichment in the full analysis and after exclusion of broadly represented genes. Strong overlap is concentrated in synaptic, postsynaptic, NMDAR-proximal, and selected excitability-associated modules, whereas E1 and R13 show little or no enrichment. **(B)** Genes recurring across enriched modules. Bars show the number of modules containing each shared gene and distinguish broadly represented genes from genes retained as identity-specific features. **(C)** Schematic summary of stage-associated program architecture in human excitatory and inhibitory populations, highlighting early depletion of selected states followed by engagement of alternative synaptic, excitability, and regulatory programs in late AD. **(D)** Conceptual summary of selective neuronal remodeling across AD stage and neuronal subtype.

**Figure S11.**
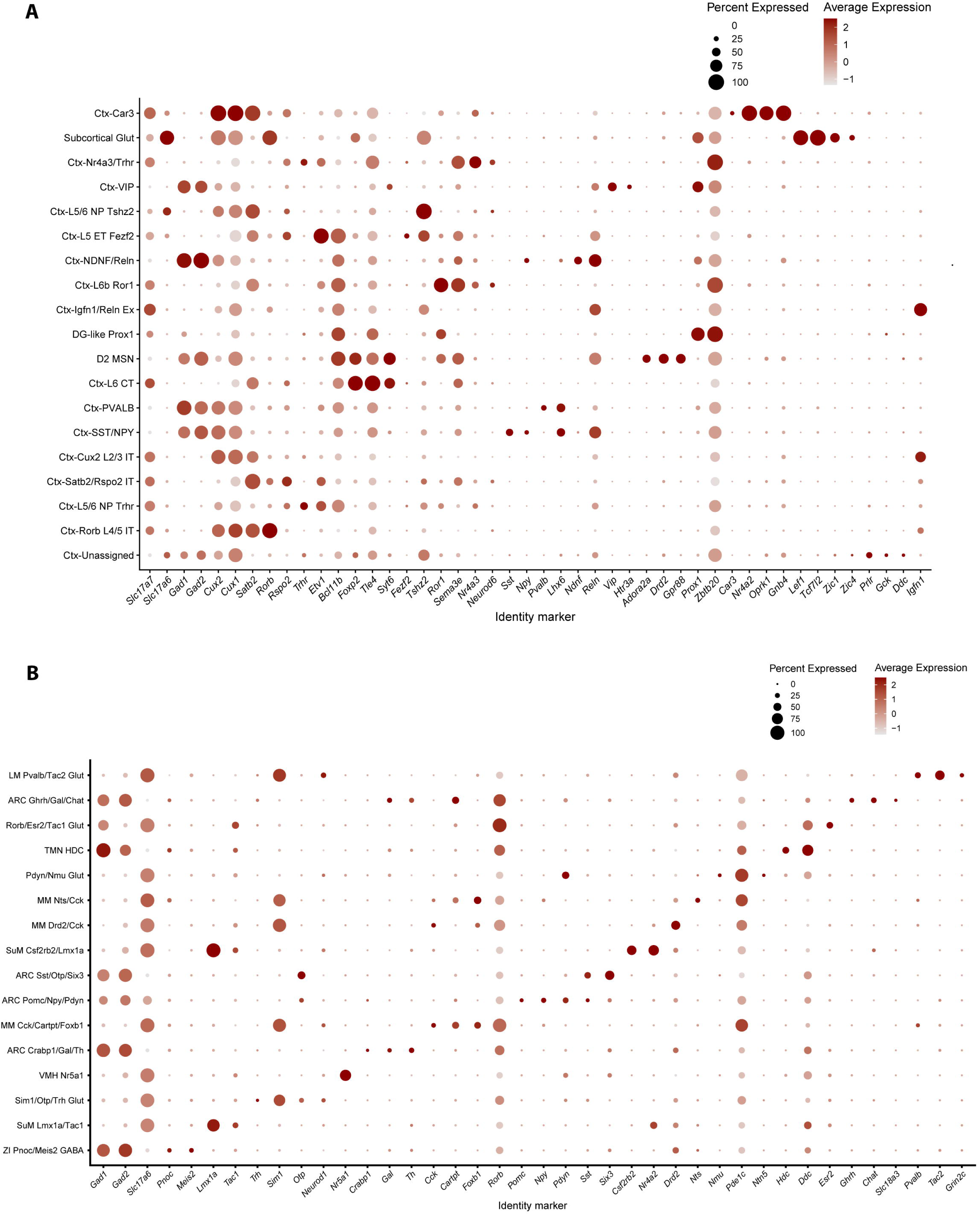
Marker-expression profiles support cortical and hypothalamic neuronal annotations in the independent TauP301S-AP cohort. **(A)** Marker-expression dot plot for cortical neuronal populations in the independent snRNA-seq dataset. Rows show annotated cortical identities and columns show selected identity markers. Dot size represents the percentage of nuclei expressing the indicated gene and color represents average expression. **(B)** Corresponding marker-expression dot plot for hypothalamic neuronal populations. Marker combinations support identities associated with arcuate, ventromedial hypothalamic, supramammillary, mammillary, zona-incerta-like, and tuberomammillary populations, including the Hdc-positive TMN histaminergic population.

**Figure S12.**
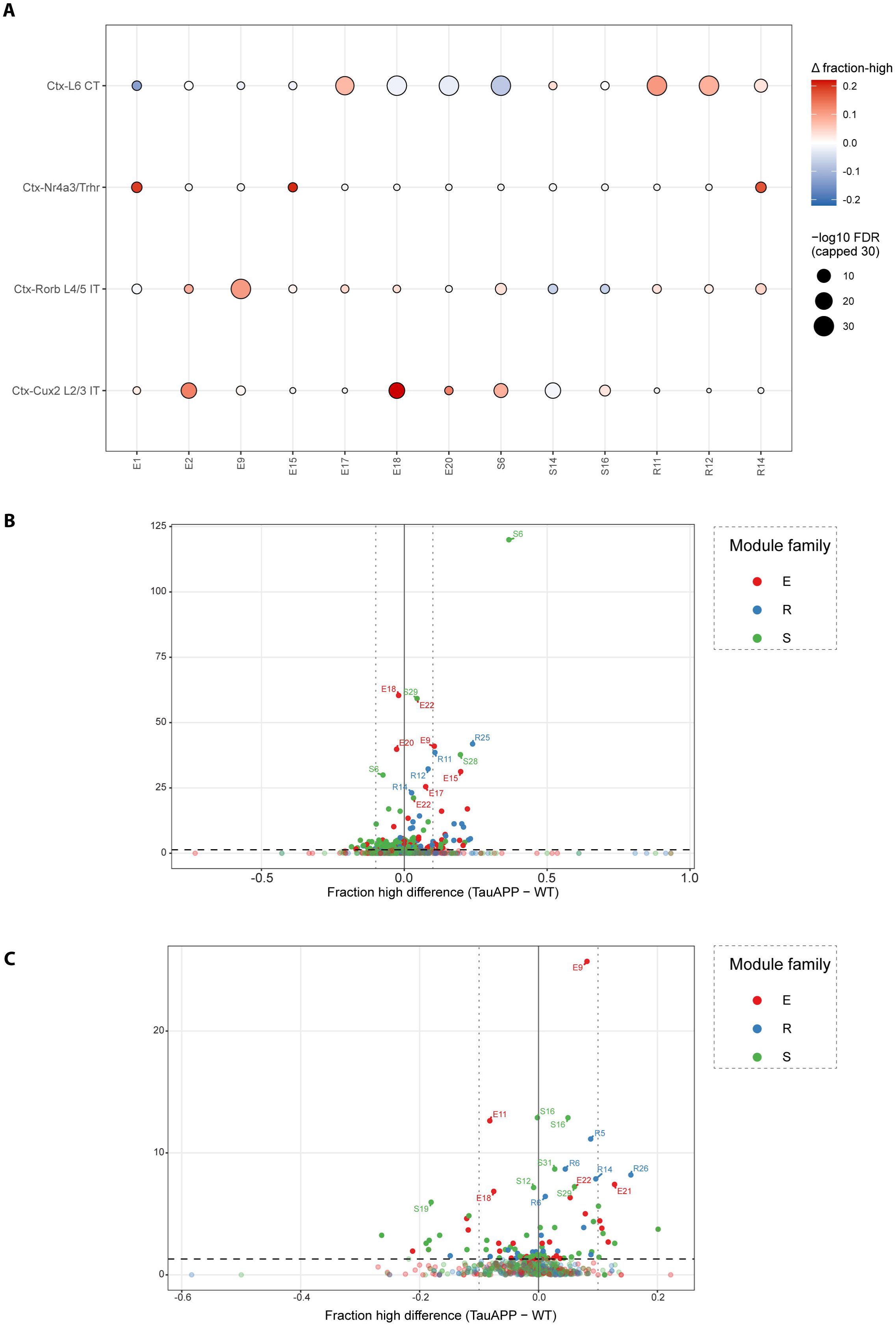
Cortical and hypothalamic neuronal populations show population-specific module remodeling in the independent TauP301S-AP cohort. **(A)** Selected module-state effects across cortical glutamatergic populations represented in both genotypes. Rows show selected E-, S-, and R-family modules and columns show Ctx-L6 CT, Ctx-Nr4a3/Trhr responsive, Ctx-Rorb L4/5 IT, and Ctx-Cux2 L2/3 IT. Circle color represents the TauP301S-AP-minus-WT difference in module-high fraction and circle size represents absolute effect magnitude, |Δfraction-high|. The four populations engage distinct combinations of excitability, synaptic, and regulatory programs. **(B)** Global distribution of module-state effects across cortical neuronal populations. Each point represents a neuronal population-module comparison. Signed Δfraction-high is used as the transcriptional-state effect size, with module family indicated by color. Vertical reference lines at Δfraction-high = -0.05 and +0.05 indicate the predefined descriptive effect-size boundaries used to identify appreciable state changes. **(C)** Corresponding distribution of module-state effects across hypothalamic neuronal populations.

**Figure S13.**
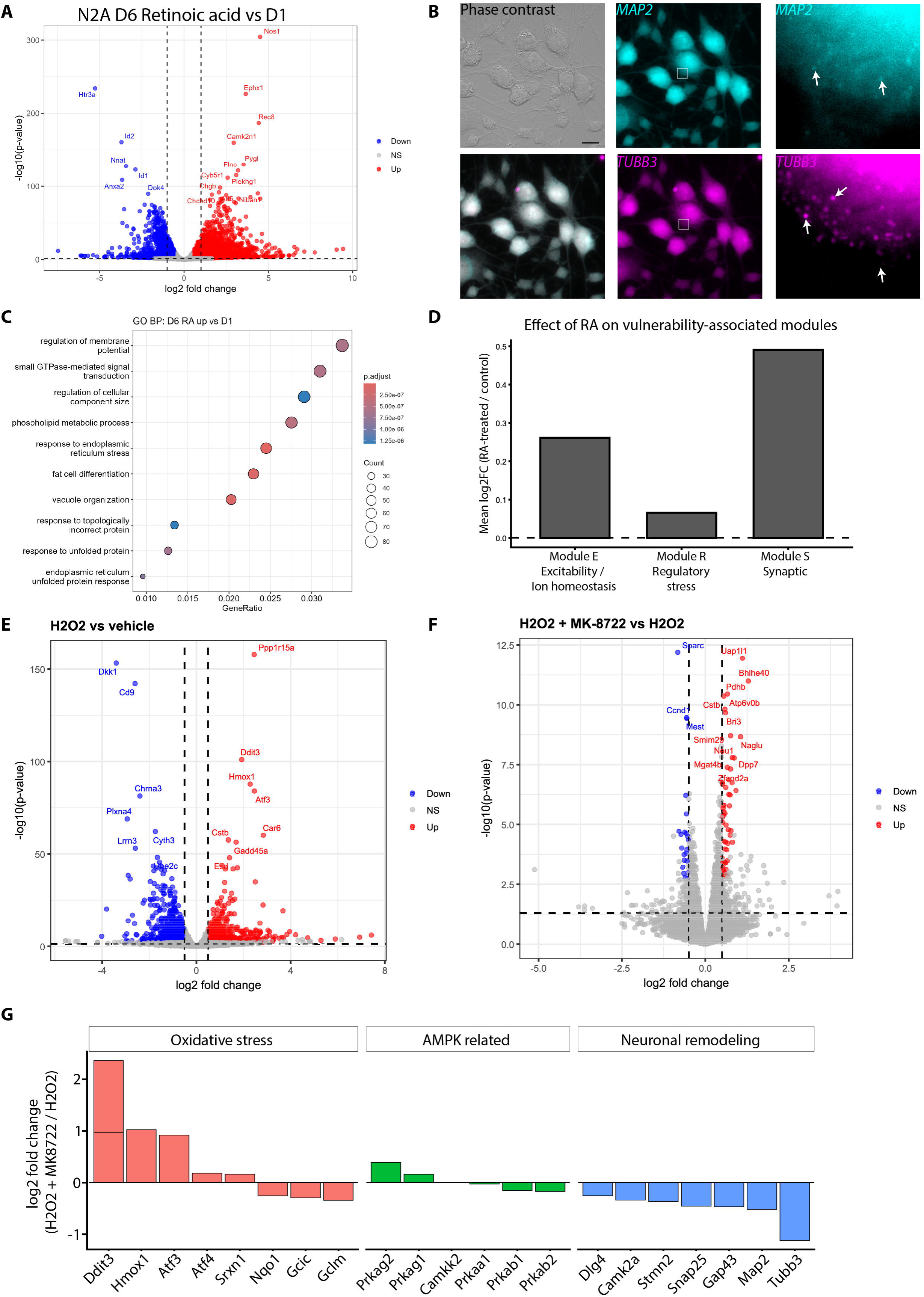
Retinoic acid conditioning and oxidative stress reveal context-dependent transcriptional responses in Neuro2a cells. **(A)** Differential-expression analysis comparing retinoic acid-conditioned Neuro2a cells at day 6 with day 1. The volcano plot shows gene-level log2 fold change and statistical support, with selected differentially expressed genes labeled. **(B)** Morphological and smFISH characterization of retinoic acid-conditioned Neuro2a cells. Phase-contrast and fluorescence images show expression of the neuronal-associated transcripts MAP2 and TUBB3. Enlarged regions illustrate transcript signal associated with cell bodies and neurite-like processes. Probe sequences and fluorophore assignments are provided in Table S19. Scale bar = 20 µm. **(C)** Gene Ontology Biological Process enrichment among genes increased following retinoic acid conditioning. Dot size denotes gene count and color denotes adjusted enrichment significance. **(D)** Effect of retinoic acid conditioning on the mouse-derived neuronal-state module families. Bars summarize mean transcriptional change across represented E-, R-, and S-family module genes. **(E)** Differential-expression analysis of H O -treated relative to vehicle-treated retinoic acid-conditioned Neuro2a cells. H O produces a broad oxidative-stress-associated transcriptional response, including induction of Ddit3, Hmox1, and Atf3. **(F)** Differential-expression analysis of H O + MK-8722 relative to H O alone. Addition of the direct AMPK activator modifies the oxidative-stress transcriptional state. **(G)** Selected gene-level effects of MK-8722 under oxidative stress, expressed as H O + MK-8722 relative to H O . Genes are grouped as oxidative-stress-associated, AMPK-related, or neuronal-remodeling-associated. Positive values indicate increased and negative values reduced expression after MK-8722 treatment in the H O context.

## Supplemental Table

**Table S1.** Neuronal population annotation and marker genes. Identity assignment and sampling depth for all annotated neuronal populations, including those excluded from composition analyses.

**Table S2.** Pathway-module definitions. The finalized E-, S-, and R-family module panel used throughout the study.

**Table S3.** Pathway-module gene membership. Gene content of each module after ortholog conversion.

**Table S4.** Detection-depth and transcript-complexity sensitivity analyses. Tests of whether the principal module-state effects are attributable to transcript recovery rather than biology.

**Table S5.** Relative neuronal representation in the Tau4RΔK-AP discovery cohort. Composition across neuronal classes and populations, with biological-sample-level statistical support.

**Table S6.** Aggregate transcriptional-state remodeling across the Tau4RΔK-AP progression cohort. Remodeling magnitude alongside relative representation for each population and age.

**Table S7.** Distributional remodeling metrics for selected neuronal states. Location, occupancy, displacement, and width of the per-cell state distributions highlighted in Figure 3.

**Table S8.** AUCell threshold-sensitivity analysis. Stability of the selected module-state effects across alternative definitions of the module-high state.

**Table S9.** Association of E1 state with age-associated changes in relative neuronal representation. Tests of whether early transcriptional state orders later representation change.

**Table S10.** RNA-ATAC neuronal population correspondence. Population matching used for cross-modal comparison.

**Table S11.** Module-associated chromatin accessibility. Accessibility-level effects alongside the corresponding transcriptional effects.

**Table S12.** Human neuronal-state remodeling across Alzheimer’s disease stages. Engagement of the mouse-derived modules across human neuronal populations and disease stages.

**Table S13.** Cross-species E1 remodeling. E1 state across mouse populations and ages and human populations and disease stages.

**Table S14.** Overlap between the mouse-derived module framework and the human Exc NRGN BEX1 identity program. Correspondence between the module panel and an independently defined depletion-associated neuronal identity.

**Table S15.** Differential gene expression in TauP301S-AP cortex. Genotype-associated expression differences by cortical neuronal population.

**Table S16.** Differential gene expression in TauP301S-AP hypothalamus. Genotype-associated expression differences by hypothalamic neuronal population.

**Table S17.** Relative representation and aggregate transcriptional-state remodeling in the TauP301S-AP cohort. Both dimensions across cortical and hypothalamic populations.

**Table S18.** Neuro2a bulk RNA-sequencing and E1 transcriptional response to oxidative stress and MK-8722. Transcriptional effects of the treatment contrasts, including the E1 response.

**Table S19.** smFISH probe and barcode information. Reagents used for smFISH in retinoic acid-conditioned Neuro2a cells.

